# Systematic dissection of phosphorylation-dependent YAP1 complex formation elucidates a key role for PTPN14 in Hippo signal integration

**DOI:** 10.1101/2022.03.13.484137

**Authors:** Federico Uliana, Rodolfo Ciuffa, Ranjan Mishra, Andrea Fossati, Fabian Frommelt, Martin Mehnert, Eivind Salmorin Birkeland, Matthias Peter, Nicolas Tapon, Ruedi Aebersold, Matthias Gstaiger

## Abstract

Cellular signaling relies on the temporal and spatial control of the formation of transient protein complexes by post-translational modifications, most notably by phosphorylation. While several computational methods have been developed to predict the functional relevance of phosphorylation sites, assessing experimentally the interdependency between protein phosphorylation and protein-protein interactions (PPIs) remains a major challenge. Here, we describe an experimental strategy to establish interdependencies between specific phosphorylation events and complex formation. This strategy is based on three main steps: (i) systematically charting the phosphorylation landscape of a target protein; (ii) assigning distinct proteoforms of the target protein to different protein complexes by electrophoretic separation of native complexes (BNPAGE) and protein/phopho correlation profiling; and (iii) genetically deleting known regulators of the target protein to identify which ones are required for given proteoforms and complexes. We applied this strategy to study phosphorylation-dependent modulation of complexes containing the transcriptional co-regulator YAP1. YAP1 is highly phosphorylated and among the most extensively connected proteins in the human interactome. It functions as the main signal integrator and effector protein of the Hippo pathway which controls organ size and tissue homeostasis. Using our workflow, we could identify several distinct YAP1 proteoforms specifically associated with physically distinct complexes and infer how their formation is affected by known Hippo pathway members. Importantly, our findings suggest that the tyrosine phosphatase PTPN14 controls the co-transcriptional activity of YAP1 by regulating its interaction with the LATS1/2 kinases. In summary, we present a powerful strategy to establish interdependencies between specific phosphorylation events and complex formation, thus contributing to the “functionalization” of phosphorylation events and by this means provide new insights into Hippo signaling.

## INTRODUCTION

Two of the central principles of cell signaling regulation are the state-specific formation and dissolution of protein assemblies and the site-specific modification (post-translational modifications - PTMs) of signaling proteins, particularly phosphorylation ^1,2^. Mass spectrometry represents the method of choice to analyze both. protein-protein interactions (PPIs) and protein phosphorylation with high throughput, dynamic range, accuracy and sensitivity^3^. While affinity purification coupled to mass spectrometry (AP-MS) has traditionally been the method of choice for the identification of PPIs, newer methods have emerged that specifically identify proximal proteins (e.g. BioID)^4^ as well as groups of proteins co-separating under native conditions, therefore suggesting protein complexes (protein correlation profiling, PCP)^5,6^. For phosphorylation, phospho-peptide enrichment strategies have compensated for the frequently substoichiometric nature of these peptides, and state-of-the art efforts are routinely capable of quantifying thousands of different sites^7^. However, protein phosphorylation and protein interactions are not independent events, but rather represent two, frequently causally interdependent aspects of the same regulatory system. In most signaling studies these two aspects are dealt with in distinct experimental and computational settings, hence separating two key facets of the cellular regulatory networks. The integration of the ensuing results can indicate statistical associations between phosphorylation patterns and PPIs but they fail to establish a causal link between phosphorylation and PPIs^8,9^. Defining dependencies between phosphosites and specific interactions is limited by several technical and conceptual factors. First, the consistent and quantitative detection of phosphosites is limited by their low abundance and difficulties associated with the correct localization of the phosphate ester groups to specific amino acid residues^10,11^. Second, PPI data generated by AP-MS or proximity labeling of a bait protein indicate the identity of interacting or proximal proteins in the tested cellular context. Nevertheless, these methods fail to probe the actual composition of specific protein complexes as a function of the cell’s signaling state or to resolve the association of (phosphorylation) proteoforms with specific complexes^12^. In order to go beyond correlation, studies need to experimentally and computationally integrate accurate, deep quantification of phosphosites with spatially resolved determination of PPIs.

Intersecting literature information about human PPIs with the most recent survey of functional phosphosites identifies a small subset of proteins that are both signaling hubs and strongly regulated at the post-translational level. YAP1 stands out as a unique example because it (i) is a promiscuous interactor (top 1% in terms of number of known interactors)^13^; (ii) carries a high number of identified phosphorylation events and functional phosphosites^14,15^ (only second to p53 among top 1% promiscuous interactors); and (iii) has a well-characterized signaling role. For these reasons, we chose YAP1 in our study as a model to establish and apply a robust workflow to determine the context-specific interdependencies between phosphorylation and PPI formation (Figure S1a). Further, YAP1 is best known as a main effector of the Hippo pathway, a conserved signaling cascade that regulates tissue homeostasis and organ size. The core of the Hippo pathway is a kinase module of the Mammalian STE20-like 1/2 (MST 1/2) and Large tumor suppressor 1/2 (LATS1/2), kinases that target YAP1, and a second transcriptional co-activator, TAZ. Once the pathway is activated, MST1/2 phosphorylates LATS1/2, thus promoting activation of the kinase and consequent YAP1 phosphorylation. Phosphorylation events in YAP1 reduce its nuclear localization and binding to the TEAD family of transcription factors, thereby blocking the transcription of genes involved in cell proliferation, apoptosis and differentiation^16,17^.

Our current knowledge on the role of YAP1 phosphorylation in Hippo signaling is largely based on a limited set of widely available phosphorylation-specific antibodies. For instance, *Cell Signaling Technologies* reports antibodies for only 4 of 52 known and annotated sites^15^. However, protein phosphorylation databases suggest a significant number of additional YAP1 phosphorylation sites which at present are functionally unexplored. This bias is well exemplified by the site S127, that makes up about 50% of all low-throughput studies entry in the *Phosphositeplus* repository, but only about 10% of the high-throughput studies entries^15,18^ (Figure S1b).

In this work, we develop an integrated multi-layered proteomic workflow to study the interdependencies between protein phosphorylation and PPIs (Figure 1). In a first step, we combine cellular phosphatase inhibition with AP-MS to comprehensively map the extent and plasticity of YAP1 phosphorylation in treated and untreated cells and to determine impact of phosphorylation on YAP1 protein interactions. In a second step we separate affinity purified YAP1 complexes by Blue Native PAGE (AP-BNPAGE) from cells at different states and characterize by MS the composition of different YAP1 modules and phosphorylation state of the constituent proteins (we use here the term ‘module’ to refer to a group of co-migrating proteins, and ‘complex’ to refer to physically stable assemblies). Finally, we apply targeted proteomics to immuno-affinity purified, endogenous YAP1 complexes to quantify phosphorylation sites and interactors detected in the previous steps in a panel of cell lines with genetic deletion of Hippo pathway members previously linked to the regulation of YAP1 activity. Importantly, while the results confirm prior knowledge, they provide new molecular understanding of the impact of LATS1/2, RHOA and NF2 on YAP1 regulation, and identify the non-receptor tyrosine phosphatase 14 (PTPN14) as an important non-canonical regulator of YAP1 function. Indeed, our data show that formation of a YAP1-LATS1/2 complex and subsequent YAP1 phosphorylation requires the presence of PTPN14. We thus propose a model where PTPN14 controls YAP1 activity by facilitating LATS-YAP1 complex formation and subsequent LATS-dependent phosphorylation which, in turn, controls YAP1 complex organization in the nucleus, cytoplasm and at cell junctions. In summary, we establish generic method that systematically dissects phosphorylation-dependent complex formation as a promising avenue to understand signaling mechanisms of proteins that similar to YAP1, act as key integrator and effectors of diverse signaling inputs.

**Figure 1.**
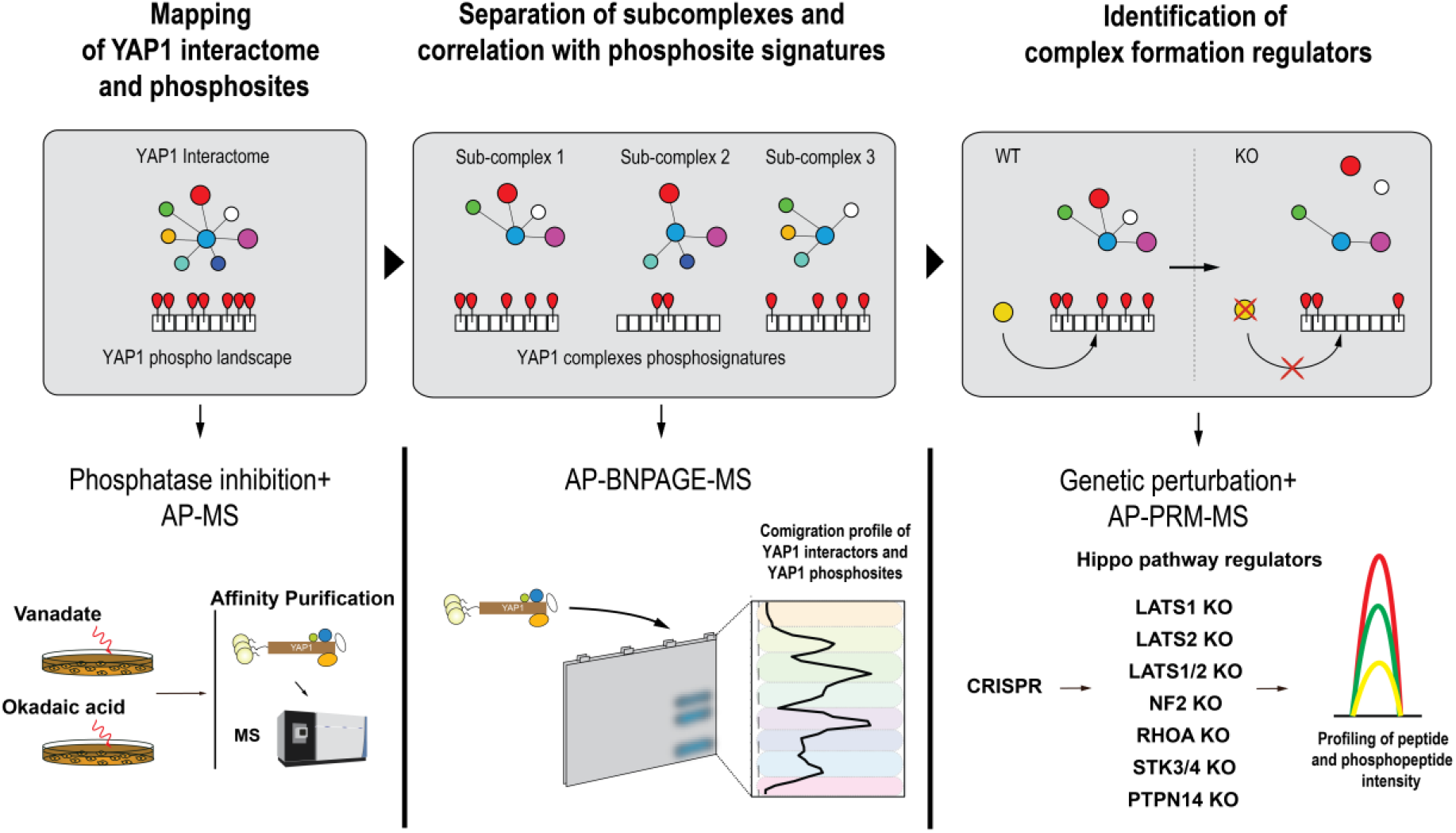
Study design. Systematic dissection of phosphorylation-dependent YAP1 complex re-organization. First, YAP1 interactors and phosphosites were identified and quantified in steady-state and upon perturbation with phosphatase inhibitors (left). In a second step, YAP1 interactors were separated on a BNPAGE and physically distinct modules and the associated YAP1 proteoforms were analyzed by mass spectrometry (center). Finally, using an exhaustive mapping of the endogenous interactome and phosphoproteome of YAP1 as a reference, the effect of a panel of genetic deletions on both levels has been measured (right).

## RESULTS

### Plasticity of the phosphorylation-dependent YAP1 interactome

To comprehensively map the extent and plasticity of YAP1 phosphorylation and its role in shaping the interactome of YAP1, we performed affinity purification and mass spectrometry (AP-MS) to quantify YAP1 interactors and phosphorylation sites in response to phosphatase inhibition (Figure 2a, Figure S2a/b/c/d/e/f/g/h/i and Supplementary Table 1). Specifically, we used SH-tagged YAP1 ectopically expressed in HEK293 cells under the control of a doxycycline-inducible promoter. We performed triplicate measurements at two time points (2, 20 minutes) after treatment of cells with the tyrosine phosphatase inhibitor vanadate, and at two time points (60 and 150 minutes) after treatment with okadaic acid, a serine/threonine phosphatase inhibitor^19^. After stringent data filtering using a SAINT probability^20^ >0.90 for interactors assignment and a PTM localization score > 0.8 for phosphosites (see Materials and Methods for details), we mapped 25 YAP1 phosphorylation sites (Figure 2b., left) and detected 32 high confidence interacting proteins (Figure 2b., right). Remarkably, 96% of the claimed phosphosites and 84 % of the identified interactors are supported by published evidence and corroborates the precision and reliability of the presented information.

**Figure 2.**
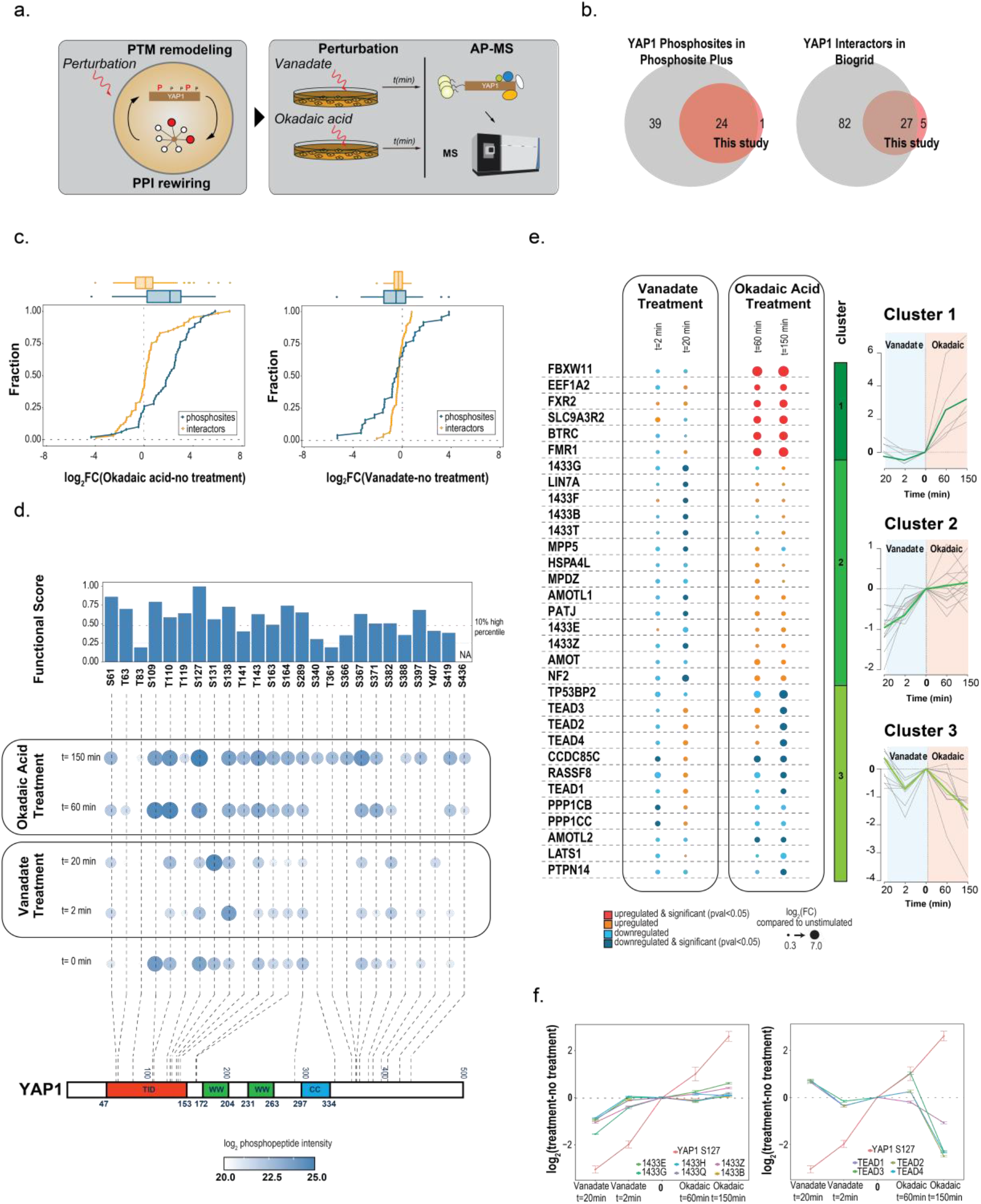
Plasticity of the phosphorylation-dependent YAP1 interactome. **a.** AP-MS approach to profile YAP1 phosphorylation changes and interactome rewiring after phosphatase inhibitor treatment. **b.** Overlap of identified and annotated phosphosites (left) and interactors (right) between this study and reference databases. Phosphosites annotated in *Phosphositeplus* (CST) and YAP1 interactor annotated in BioGRID as “physical and direct interactor” in at least two independent experiments were considered in the reference databases **c.** Empirical cumulative density function (ECDF plot) for YAP1 interactors and phosphosites after stimulation with okadaic acid (left) and vanadate (right). x axis represents the log2FC for the respective perturbation versus the control sample (untreated). **d.** Kinetics of YAP1 phosphorylation sites. After treatment with vanadate (2, 20 minutes) and okadaic acid (60, 150 minutes), YAP1 phosphosite abundance was measured by MS after YAP1 AP-MS. Size and color of circles represent the average abundance of phosphopeptides normalized for YAP1 intensity. Barplot on top indicates the functional score associated with each site as reported in Ochoa et al., 2019. On the bottom, phosphosites are localized onto YAP1 primary sequence (lower part). **e.** Kinetics of YAP1 interactors upon phosphatase treatment. After treatment with vanadate (2, 20 minutes) and okadaic acid (60, 150 minutes), high confidence YAP1 interactors (SAINT SP>0.9) were profiled by MS after YAP1 AP-MS. The dot size represents log2 fold change from triplicate experiments compared to non-treated samples. Interactors are fuzzy-clustered based on the kinetic profile of fold change compared to no treatment condition (right panel). **f.** Fold change profile of YAP1 S127 phosphosite with 14-3-3 protein family (left) and with TEADs protein family (right).

Okadaic acid and vanadate differentially affected the direction and magnitude of YAP1 phosphorylation and, to a lesser extent, interactor association (Figure 2c). Cumulative density function shows that okadaic acid had an impact on a larger number of phosphosites and caused a dramatic alteration, up to 100fold, of several PPIs, while vanadate affected a lower number of phosphosites and protein interactions (Figure 2c). As expected, the use of phosphatase inhibitors improved phosphosite detection: only 15 YAP1 phosphosites were detected without treatment, whereas 25 phosphosites were detected cumulatively after the addition of the respective phosphatase inhibitors (Figure S2f.). Identified YAP1 phosphosites are primarily localized in the N-terminal TEAD interaction domain (aa 47-153, TID) or at the C-terminus (aa 335-500) of YAP1 (Figure 2d). The former region is characterized by phosphosites that have a higher functional score^14^ than the latter. Interestingly, most sites, regardless of the sequence location, show very high functional scores (top 10% percentile or higher) against the entire human phosphoproteome^14^ (Figure 2d, top).

Next, we studied whether the temporal response to vanadate and okadaic acid would reveal distinct changes in YAP1 interactors. To do that, we used a fuzzy clustering approach to group protein response profiles, considering both treatments (see material and methods). Rewiring of the YAP1 interactome under these perturbation conditions indicated three main clusters of alteration (Figure 2e, Figure S2i). These clusters display distinct association dynamics and suggest that proteins exhibiting similar behavior may be part of the same complexes. Indeed, several structurally or functionally related proteins clustered together under the conditions tested, indicating that their interaction with YAP1 is modulated coordinately and controlled by YAP1 phosphorylation status. The first cluster contains the F-box proteins BTRC and FBXW11, which interact more strongly with YAP1 after okadaic acid treatment compared to untreated cells. BTRC and FBXW11 are known to mediate SCF-dependent ubiquitination and degradation of YAP1, after phosphorylation of S397, S400 (not detected) and S403 (not detected)^21^. The second cluster consists of apicobasal polarity proteins (AMOT, INADL, MPDZ, MPP5, NF2) and 14-3-3 proteins. These proteins interacted less strongly with YAP1 in presence of vanadate compared to untreated cells. The third cluster consisted of proteins that showed reduced binding to YAP1 in the presence of okadaic acid compared to untreated cells. It includes the members of the ASPP/PP1A complex (CCDC85C, TP53BP2, RASFF8, PPP1CB and PPP1CC) and TEAD protein family members. Consistent with previous reports, we found an inverse YAP1 association behavior of TEAD compared to 14-3-3 proteins, whereby, after okadaic acid treatment, the hyperphosphorylation of YAP1 in the TEAD interaction domain (TID) (S127) resulted in a strongly reduced binding of YAP1 for TEAD proteins (TEAD1,2,3,4). In contrast, reduced phosphorylation of YAP1 S127 correlated with a strong decrease in binding of 14-3-3 proteins (1433F, 1433B, 1433T, 1433E, 1433Z) (Figure 2f). This is consistent with the previous finding that pS127 acts as a docking site for 14-3-3 proteins ^22^ and causes YAP1 translocation. Taken together, these data provides an extensive, unbiased map of YAP1 phosphosites and their responsiveness upon phosphatase inhibition. Further, it indicates how changes in the phosphorylation state of YAP1 are correlated with an organized reshaping of its interactome around three functionally coherent clusters of proteins with distinct association dynamics.

### Deconvolution of native YAP1 complexes by protein correlation profiling

Since the AP-MS data represents binary interactions resulting from the sum of concurrently purified YAP1 complexes, AP-MS data does not inform about the presence of differentially phosphorylated YAP1 complexes in the same sample^23^. To assign the identified YAP1 interactors to specific YAP1 subcomplexes we subjected an affinity purified YAP1 complex mixture to electrophoretic native size fractionation (BNPAGE)^24^(Figure 3a). Specifically, we separated YAP1 complexes along the axis of native electrophoretic separation (molecular weight), excised 64 consecutive gel slices and used untargeted MS to measure the abundance of YAP1 phosphopeptides and interactors (identified in the experiment above, Figure S2h) to generate migration profiles of protein and phosphosites (Figure 3b and Figure S3, S4, S5a/b/c/d/e/f; Supplementary Table 2). Migration profiles of the same entity (protein, phosphosites) showing multiple peaks (i.e. potentially being present in multiple assemblies) were deconvoluted based on the detection of local maxima, and the resulting single peaks from different proteins/phosphosites were grouped by unsupervised hierarchical clustering into co-migrating modules (see Material and Methods and the reference^25^). Critically, the YAP1 profile across the analyzed fractions indicates the existence of electrophoretically well-resolved peaks of varying abundance and MW (Figure S5d and S3, S4 for the visualization of raw and smoothed profiles). Strikingly, both, YAP1 phosphosites and interactors exhibit similarly discrete partitioning across the fractionation dimension (Figure 3b/c and Figure S3, S4). Of note, the BNPAGE protocol does not affect the original overall abundance range and stoichiometries of YAP1 interactors, as their relative abundances are highly correlated with the unseparated YAP1 interactome (Figure 3d). Migration profile analysis indicated separation of YAP1 modules into nine distinct mobility clusters with specific YAP1 phosphorylation patterns (Figure 3b). Several lines of evidence support the notion that these modules are indeed biologically relevant entities and not the result of coincidental co-migration. When compared with random subsets of YAP1 interactors (see Material and Methods), proteins belonging to the same clusters were (i) significantly more often found to interact with one another (BioGRID)^13^ (Figure 3e); (ii) significantly more strongly associated with the same GO cellular compartment (Figure 3f); (iii) more frequently co-regulated upon phosphatase inhibition (Figure 3g; Figure 2e); (iv) and more likely to be part of known, stable complexes (ASPP-PP1, RICH1-AMOT, Figure 3c) or to contain homologous proteins (e.g. TEAD, 14-3-3) (Figure 3c).

**Figure 3.**
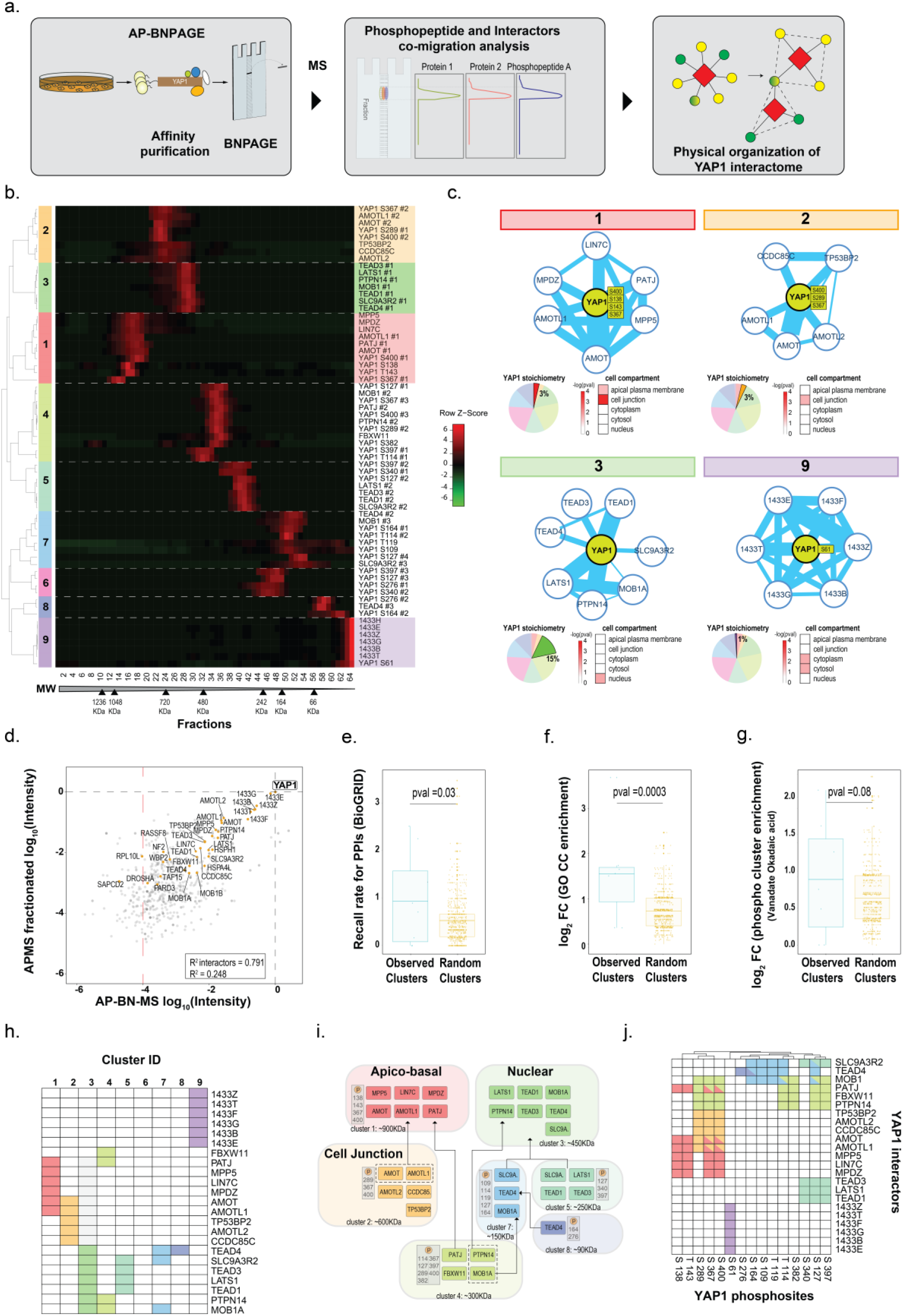
Deconvolution of native YAP1 complexes by protein correlation profiling. **a.** Workflow of AP-MS combined with BNPAGE to investigate the organization of YAP1 interactors and phosphosites. After native elution, YAP1 interactors and YAP1 proteoforms were fractionated based on their electrophoretic mobility under native conditions. Quantitative proteomics data was obtained from the integration of MS1 signal over 64 gel fractions to generate migration profiles of proteins and phosphosites. **b.** Unsupervised hierarchical clustering of YAP1 interactors and phosphosites intensity profiles. **c.** Composition, YAP1 phospho-signature, stoichiometry (pie chart) and localization (heatmap) of 4 selected modules (1,2,3,9). Edge thickness corresponds to the number of physical PPIs annotated in BioGRID database. **d.** Protein intensity correlation between YAP1 AP-MS and AP-BNPAGE (provided by the total intensity sum of all measured fractions). R^2^ for reported YAP1 interactors (0.791) and background (0.248) is reported in the lower box. **e./f./g.** Co-migrating proteins isolated by BNPAGE display a higher degree of relatedness when compared with randomized clusters, as measured based on reported PPI (BioGRID) (**e.**), associated GO cellular component terms (“apical plasma membrane”, “cell junction”, “cytoplasm”, “cytosol”, “nucleus”)(**f.**), and kinetics of YAP1 binding upon phosphatase inhibition (**g.**). In all cases, observed clusters were compared against randomized clusters generated by sampling 1000 times subsets of the identified YAP1 interactors with the same size as the observed clusters. The boundaries of the box plot correspond to the quantiles Q1 (25%) and Q3 (75%). Lower and upper whiskers are defined by Q1 −1.5IQR and Q3+ 1.5IQR. **h.** Composition of the 9 identified modules (group of co-migrating proteins). **i.** Graphical representation of modules identified by AP-BNPAGE experiment. Each module is characterized by PTM signature and the estimated molecular weight from AP-BNPAGE experiment. Relationship between modules (fragment or assembly) are indicated by arrows **j**. Graphical representation of the identified modules and the comigrating YAP1 phosphosites based on the clustering assignment.

We next examined the distribution of the YAP1 interactors identified above across the nine identified modules (Figure 3h). We found three high molecular weight modules (modules 1-3), several partially overlapping modules of intermediate size (modules 4-8) and smaller molecular weight YAP1 complexes containing 14-3-3 proteins (module 9). The first module contains mostly apical-basal proteins, including PATJ, MPP5, LIN7C, MPDZ, AMOT and AMOTL1 of the RICH1/AMOT polarity complex^26^. The second module encompassed tight junction proteins, including members of the ASPP/PP1 complex^27^ involved in YAP1 S127 dephosphorylation^28^, in addition to AMOT, AMOTL1 and AMOTL2. The third module primarily consisted of known nuclear interactors of YAP1, including the TEAD transcription factors TEAD1, TEAD3, TEAD4, LATS1 and its activator MOB1 as well as PTPN14. Modules 5, 7 and 8 are most likely fragments or assembly intermediates of this nuclear module, while we interpret module 4 as a convolution of a fragment of the nuclear module and two additional proteins (Figure 3i). Overall, our profile analysis separates YAP1 the interactome in distinct complexes that are linked to its signal integration and effector function.

In most modules we were able to identify specific YAP1 phospho-signatures (ensemble of phosphosites) (Figure 3c, 3j). For example, in the apical cell polarity complex (module 1) YAP1 was phosphorylated on S138, S143, S367 and S400. Among these, S138 and S367 are phosphorylated by CDK1 through a mechanism involving the interaction with the polarity protein PATJ^29^. This is in striking contrast to the fragments or assembly intermediates of the nuclear module (modules 4, 5, 7, 8, Figure 3i) where YAP1 is richly phosphorylated on several sites and the larger nuclear module itself (module 3), where no YAP1 phosphorylation sites were detected. This pattern cannot be explained by the overall abundance of YAP1 in these different complexes, since YAP1 intensity is comparable in the nuclear module and in some of the submodules (Figure S5d). These results suggest that S127 may not be the sole regulator of nuclear/cytoplasmic transport, but that it may require dephosphorylation of multiple sites. Overall, our strategy shows that integrated MS-based analysis of complex composition and phosphorylation state, combined with native fractionation of purified complexes, can deconvolute the interactome (sum of all binary interactions) in biologically meaningful YAP1 complexes and assign complex-specific YAP1 proteoforms to them.

### Identification of YAP1 phosphorylation and interactors recapitulate known and suggest new control mechanisms

We have thus far mapped the effects of phosphorylation changes on the interactome of total YAP1 and further resolved the YAP1 interactome in co-migrating protein groups (complexes), of which each was associated with a specific YAP1 phospho-signature/proteoforms. To relate the observed YAP1 proteoforms to the regulation of the Hippo pathway we analyzed YAP1 phosphorylation and complex formation in a panel of HEK293A knock out (KO) cell lines where each cell line lacked a key regulator of the Hippo pathway (Figure 4a)^30^. We first established a protocol to affinity-purify endogenous YAP1 with custom-generated anti-YAP1 antibodies. Compared to the inducible ectopic expression of YAP1 used in the previous experiments (Figure 2), this approach is more compatible with systematic YAP1 interactor analysis in HEK293A mutants lacking critical Hippo signaling components and more reliably reflects endogenous stoichiometries. To maximize the specificity of our purification, we used a double control strategy by using the flow-through of the antibody purification as non-specific antibody (aB control) and a YAP1 KO HEK293A line (cell line control) to control for variation of expression following the genetic perturbation (see material and methods for details) (Figure 4b and S6a/b/c/d/ f/ and Supplementary Table 3). Importantly, we could recover almost all interactors previously defined with AP-MS of fractionated ectopically expressed YAP1 with high specificity and sensitivity (AUC 0.88 and 0.85) (Figure S6e, S6g). A global description of all YAP1 interactors and phosphosites identified in all experiments of this work is reported (Figure S8f/g, respectively for interactors and phosphosites).

**Figure 4.**
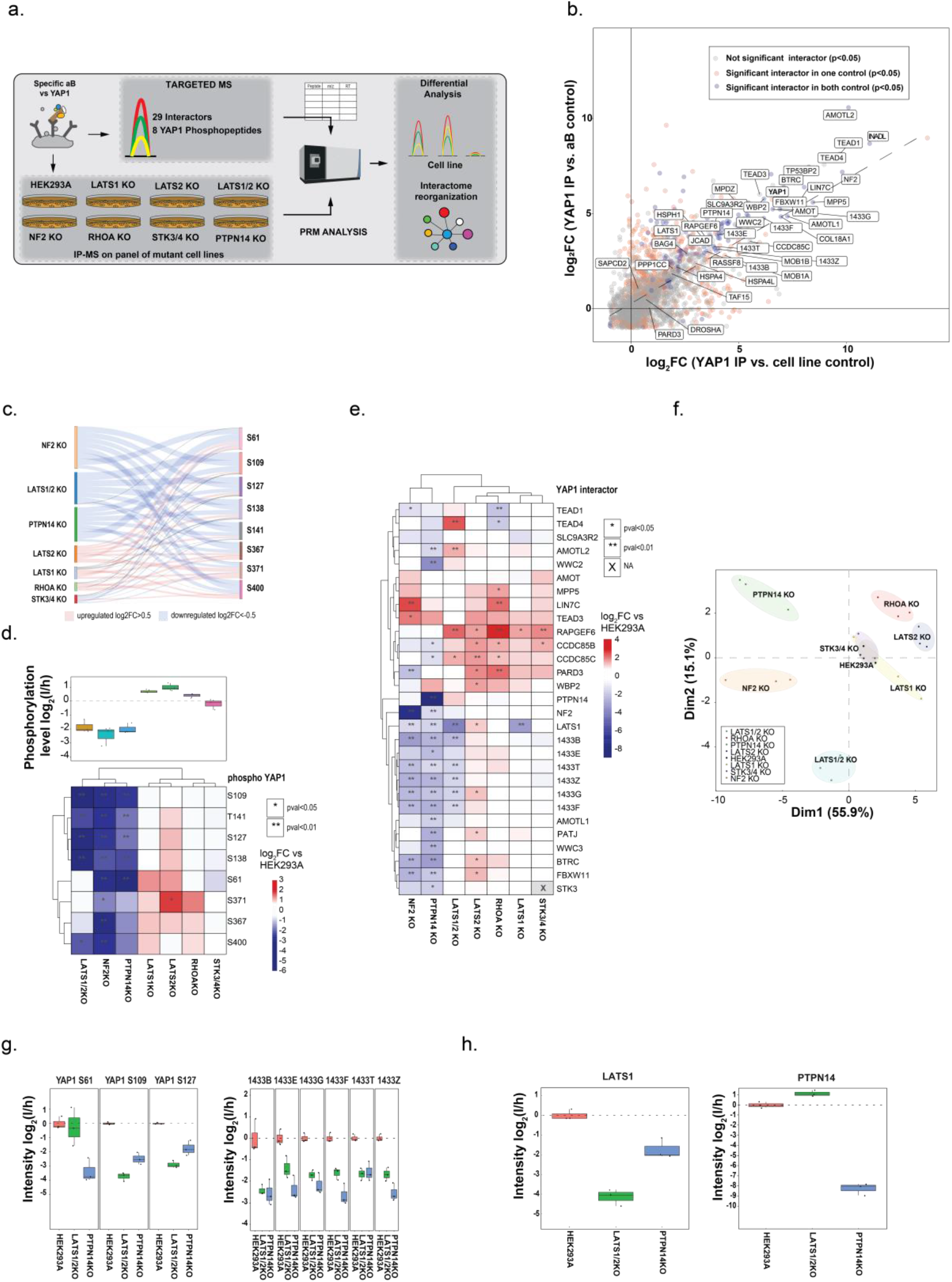
Identification of YAP1 phosphorylation and interactors recapitulate known and suggest new control mechanisms. **a.** Experimental workflow for profiling endogenous YAP1 phosphorylation and interaction changes in cells lacking known Hippo pathway members. After YAP1 endogenous immuno-affinity purification from the indicated mutant HEK293A cells, phosphopeptides and interactors were quantified by targeted proteomics. **b.** Proteins enriched in the YAP1 endogenous immune-affinity purification with two different controls cell line control and aB control. Cell line control is performed with YAP1 IP-MS from YAP1 KO cells and aB control with non-specific control antibody IP-MS from HEK293A. Proteins identified and filtered as interactors (SP>0.9) in fractionated AP-MS from HEK293A cells expressing epitope tagged YAP1 are annotated. Protein significantly enriched in both controls are marked in blue, in orange those significantly enriched with only one control, in grey those not significantly enriched. **c.** Sankey plots shows the effect of protein deletion on YAP1 phospholandscape. Color code indicates an increase compared to wild type (log2>0.5, red) or decrease (log2<-0.5, blue). **d.** Unsupervised hierarchical cluster of YAP1 phosphopeptides in a panel of seven cell lines with genetic deletions of indicated Hippo signaling genes. Values reported in the heatmap represent the log2 fold change of phosphopeptide intensity average from three biological replicates compared to parental cell (HEK293A). Upregulated and downregulated phospho- peptides are shown in red and blue respectively; significant changes are marked with asterisks. On the top, boxplot shows the average phosphorylation level of 8 monitored YAP1 phosphopeptides per condition (n =3). The boundaries of the box plot correspond to the quantiles Q1 (25%) and Q3 (75%). Lower and upper whiskers are defined by Q1−1.5IQR and Q3+ 1.5IQR. **e.** Unsupervised hierarchical cluster of YAP1 interactors in a panel of seven cell lines with Hippo genetic deletions. Values reported in the heatmap represent the log2 fold change of YAP1 interactor intensity average from three biological replicates compared to parental cell (HEK293A). Upregulated and downregulated phospho-peptides are shown in red and blue respectively; significant changes are marked with asterisks. **f.** Principal component analysis based on both phosphorylation and interaction data. Various mutants are highlighted in different colors and every dot represents a replicate. **g.** Intensities (log2) of identified YAP1 phosphopeptides with LATS1 sequence motif (S61, S109, S127) (left) and 14-3-3 protein family in the indicates cell lines (n=3). The boundaries of the box plot correspond to the quantiles Q1 (25%) and Q3 (75%). Lower and upper whiskers are defined by Q1 −1.5IQR and Q3+ 1.5IQR. **h.** Intensities of PTPN14 and LATS1 after YAP1 immuno-affinity purification in the indicated cell lines(n=3). The boundaries of the box plot correspond to the quantiles Q1 (25%) and Q3 (75%). Lower and upper whiskers are defined by Q1 −1.5IQR and Q3+ 1.5IQR.

Next, we used targeted mass spectrometry to analyze a panel of KO cell lines lacking key component of the Hippo pathway^30^, including the kinases LATS1, LATS2, LATS1/2, STK3/STK4, the GTPase RHOA, NF2, the phosphatase PTPN14 and YAP1 itself as a control. Gene deletion were confirmed by the absence of the respective proteins as measured by targeted proteomics and western blot (only PTPN14KO, Figure S7e), except for STK3/4 KO and RHOA KO cells which showed about 15% and 45% of residual STK4 and RHOA levels compared to parental controls, respectively (Figure S7a/b/c/d and Supplementary Table 4). Importantly, protein expression of other Hippo-pathway regulators was only mildly affected by the gene deletions (Figure S7c), suggesting that disruption of the Hippo network did not significantly alter protein expression or stability.

Finally, we performed endogenous YAP1 purifications in triplicate for each cell line, followed by targeted PRM measurements using heavy labelled reference peptides. Overall, we quantified 29 interacting proteins and 8 YAP1 phosphopeptides (Figure S8a/b/c/d/e; Supplementary Table 5 for a summary of all YAP1 peptides monitored in the targeted experiments). To interpret the acquired data, we performed unsupervised hierarchical clustering separately on phosphopeptide intensities (Figure 4c) and protein intensities (Figure 4e). YAP1 phosphopeptide values were normalized to YAP1 protein intensity to differentiate variations in protein abundance from changes in phosphorylation.

Sankey plot (Figure 4c) and the average phosphorylation levels (Figure 4d, upper panel) show that phosphorylation levels of YAP1 clearly separated a group of mutants that caused YAP1 hypophosphorylation consisting of NF2, LATS1/2, and PTPN14 KO cells from a group showing a mild increase in phosphorylation, consisting of the LATS1, LATS2, RHOA KO cells. In contrast, the STK3/4 double mutant cells only showed a very moderate effect on YAP1 phosphorylation on the tested sites ^30^ (Figure 4c/d). Among the tested phosphopeptides, two distinct clusters with somewhat complementary behaviors were observed. The first cluster consisted of sites located N-terminally, specifically S109; S127; S138; S143, consistently showing a highly significant dephosphorylation in the LATS1/2, NF2 and PTPN14 mutants, and no or mild upregulation in the LATS1, LATS2 and RHOA mutants. The second cluster consisted of sites located C-terminally, specifically sites S371; S379; S400, showing weaker downregulation in the LATS1/2, NF2 and PTPN14 mutants and stronger upregulation in LATS1, LATS2 and RHOA mutants. Site S61 displayed a more complex modulation, as shown in Figure 4d. Although the peptide encompassing S61 contains a LATS consensus motif ^31^, phosphorylation of this site is not affected by the deletion of LATS1/2, implying a role for other kinases, as already suggested by *in vitro* studies^32–34^ .

Unsupervised hierarchical clustering of YAP1 interactor intensities closely mirrored the clustering of phosphopeptides. The LATS1/2, PTPTN14 and NF2 KO cells showed a systematically decreased interaction with cytoplasmic proteins (14-3-3 proteins). In contrast, the others mutants (LATS1, LATS2, RHOA, STK3/4) revealed an increased or unchanged association, as indicated by the stable profile of 14-3-3 proteins (Figure 4e). In several respects, these data are in agreement with previous knowledge: our analysis confirms the positive role on YAP1 phosphorylation by NF2 and LATS1/2 already observed by others^16,35,36^, but adds a quantitative phosphosite-level resolution absent in previous analyses^30^. As reported previously, deletion of RHOA (although not quantitative), which mediates the mechanical stress-induced activation of YAP1, induces YAP1 hyper-phosphorylation and reduces its nuclear localization^30,37^ (Figure 4c/d/e). Finally, LATS2 KO cells and, to a lesser extent, LATS1 KO cells resulted in mild, but widespread hyperphosphorylation of YAP1 sites, in contrast to the strong downregulation driven by the double mutant. These results confirm that the two kinases are redundant, corroborating the characterization of the KO from prior phospho-tag experiments^30^. We surmised that the upregulation of the phosphopeptides in the single KO mutants could be due to a compensatory mechanism, whereby the loss of one kinase leads to increased expression of the other. Analysis of the levels of LATS2 in LATS1 KO lysates supports this hypothesis (Figure S7f).

Remarkably, we observed that YAP1 phosphorylation and complex formation patterns in the absence of the phosphatase PTPN14 closely resembles those in NF2 and LATS1/2 KO cells. This observation was further confirmed by a principal component analysis on the combined interactome and phosphoproteome data (Figure 4f), showing a pronounced separation of these three mutants along the major component as compared to the other mutants tested (1^st^ dimension: 55.9% of explained variance). Because YAP1 phosphorylation pattern and complex formation in PTPN14 KO cells resembles those found in cells lacking NF2, which is an upstream activator of LATS1/2, as well as in LATS1/2 double mutant cells, we hypothesized that PTPN14 may play an analogous role in activating YAP1 phosphorylation by LATS1/2. This is supported by several lines of evidence: (i) two of the LATS1/2 sites on YAP1, S109, S127^28^, are negatively regulated in PTPTN14 KO cells and, as a consequence, the interaction with 14-3-3 proteins is decreased because the docking site is eliminated (Figure 4g); (ii) PTPN14 KO reduces the binding between YAP1 and LATS1, but LATS1/2 KO does not decrease the amount of PTNP14 associated with YAP1 (Figure 4h., left and right, respectively); (iii) an interaction between PTPN14 and LATS1 has already been reported ^19,38,39^ and (iv) evidence for a role of PTPN14 in the modulation of YAP1 phosphorylation and activity have been provided^34,40,41^.

Taken together, our targeted proteomics of endogenous YAP1 immuno-purified from cells lacking Hippo pathway regulators resolved their roles in controlling YAP1 activity at the level of YAP1 phosphorylation and complex formation, and suggests a key role for PTPN14 in controlling LATS-dependent YAP1 regulation.

### Reduced LATS1/2-YAP1 complex formation, enhanced nuclear translocation and activation of YAP1 in PTPN14 mutant cells

Finally, we aimed to gain insights into the mechanism how PTPN14 could act as a positive regulator of LATS1/2 kinases for the control of YAP1. We first validated the effect of PTPN14 deletion on LATS1 activity by monitoring levels of phosphorylation of the known LATS1/2 substrate, site S127, on YAP1 by western blot. We found that PTPN14 KO reduced the level of YAP1 S127 phosphorylation to about 60% compared to the WT condition, confirming the MS results obtained with purified endogenous YAP1 (Figure 5a/ Figure 4d). Next, we compared YAP1 subcellular localization by immunofluorescence in LATS1/2 and PTPN14 KO cells. In keeping with our previous results as well as previously published data^30,34^, we found that YAP1 was localized in the cytoplasm in WT HEK293A cells, while the nuclear fraction increased upon removal of LATS1/2 and, to a lesser but still significant extent, upon PTPN14 deletion (Figure 5b). Finally, we compared the mRNA expression of CTGF and CYR61, two established YAP1 target genes in the two mutant cell lines. We found an approximately 3-fold increase in the levels of CTGF mRNA in both LATS1/2 and PTNPN14 mutants compared to parental HEK293A cells. Lack of PTPN14 was also associated with an increase of CYR61, which was even stronger than in LATS1/2 mutants (Figure 5c).

**Figure 5.**
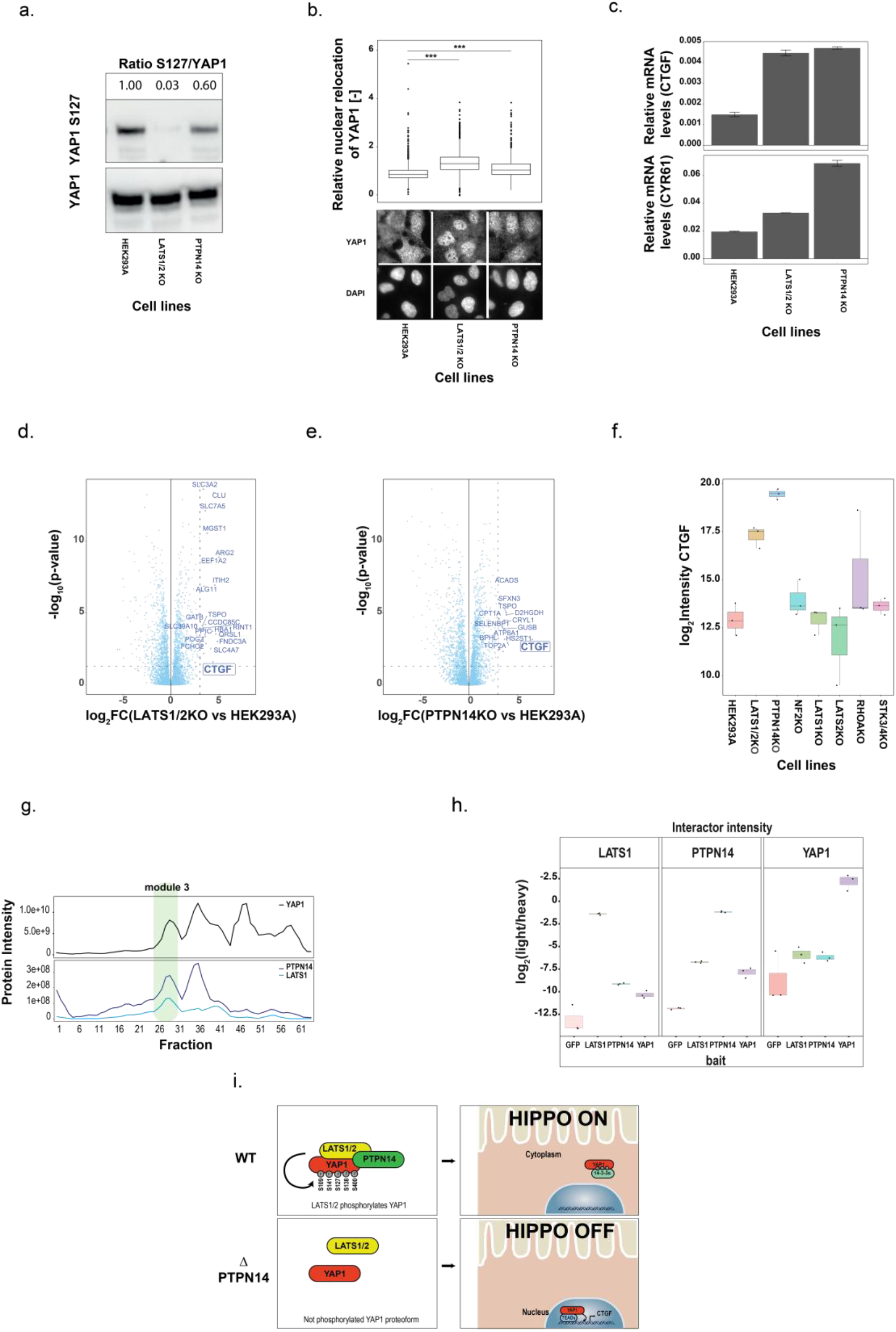
Reduced LATS1/2-YAP1 complex formation, enhanced nuclear translocation and activation of YAP1 in PTPN14 mutant cells. **a.** Immunoblot with anti YAP1 and anti phospho-YAP1(S127) antibodies on protein lysates from indicated WT and KO cell lines. Quantitative values reported above are normalized for the abundance of YAP1. **b.** Subcellular localization of YAP1 in LATS1/2 KO and PTPN14 KO cells. *Down panel:* the localization of YAP1 was probed using immunofluorescence and visualized using wide-field microscopy. The DAPI-signal was used to compare the nuclear relocation of YAP1 among the cell lines. *Upper panel:* quantification of relative nuclear relocation of YAP1 combined from three independent experiments. The boundaries of the box plot correspond to the quantiles Q1 (25%) and Q3 (75%). Lower and upper whiskers are defined by Q1 −1.5IQR and Q3+ 1.5IQR. The significance is indicated with *** for *P*<0.001 Total number of cells analyzed for HEK 293A WT, LATS1/2 KO, and PTPN14 KO cells were 1514, 1673, and 1320 respectively .**c.** CTGF and CYR61 (YAP1 target genes) transcript levels (qPCR) for HEK293A WT, LATS1/2 KO, and PTPN14 KO. Data are presented as mean values±SD. **d./e./f.** Differential protein expression data. Volcano plots displaying the log2 protein fold changes of LATS1/2 KO (**d.**) and PTPN14 KO (**e.**) compared to WT control HEK293A and the corresponding significance. Proteins with log2 FC>3 and p value <0.05 are highlighted with their gene names. **f.** Boxplot showing CTGF protein intensity level (log2) across the examined mutant cell. The boundaries of the box plot correspond to the quantiles Q1 (25%) and Q3 (75%). Lower and upper whiskers are defined by Q1 −1.5IQR and Q3+ 1.5IQR. **g.** Co-migration profile of PTPN14, LATS1 and YAP1 after YAP1 AP-BNPAGE complex fractionation. **h.** Reciprocal enrichment of the PTPN14-LATS1-YAP1 complex in the pulldowns of each of the complex members. Boxplot showing the intensity of interactors (log2) across different baits (GFP, LATS1, PTPN14, YAP1). The boundaries of the box plot correspond to the quantiles Q1 (25%) and Q3 (75%). Lower and upper whiskers are defined by Q1 −1.5IQR and Q3+ 1.5IQR. **i.** Model of the role of PTPN14: PTPN14 promoting the interaction between LATS1 and YAP1, increase phosphorylation level of YAP1 (S109, S141, S127, S138, S400) which, in turn, leads to YAP1 inactivation and cytoplasmic retention.

We next wished to validate the finding at the proteome level by performing proteome profiling across the KO cell lines using data independent MS acquisition (DIA). We identified 4436 proteins (Figure 5d/e/f, Figure S9a/b/c/d, Supplementary Table 6), and carried out differential expression analysis to identify proteins showing differential abundance under either LATS1/2 or PTPN14 KO (see Materials and Methods). Consistent with the above results, we confirmed increased CTGF protein levels in only LATS1/2 mutant cells and even greater CTGF expression in PTNPN14 KO cells (Figure 5f.).

These orthogonal lines of evidence strongly support an involvement of PTPTN14 in the regulation of LATS1/2 and YAP1 activity, but do not provide clear indications about the underlying mechanism. Because our interaction data indicates that LATS1/2 are not required for the YAP1-PTPN14 interaction, but that the reciprocal is true (Figure 4h, left and right, respectively), we propose the existence of a trimeric complex where PTPN14 mediates the interaction between YAP1 and LATS1/2 kinases. This putative complex would reminisce the characterized trimeric complex LATS-PTPN14-KIBRA^41^, in that YAP1 and KIBRA shares two WW domains with a good alignment score (BLASTp analysis, *p*=5e-16) and it is reported that YAP1 associates with PTPN14 in a WW/PPxY-dependent manner ^34,40,42,43^. The BNPAGE data supports the hypothesis for the presence of YAP1- LATS1-PTPN14 complex and indicates co-migration of the three proteins in a module with other nuclear proteins (Figure 5g; Figure 3c, module 3). To further corroborate this finding, we carried out quantitative reciprocal AP-MS in HEK293 cells expressing epitope tagged YAP1, LATS1, PTPN14 and GFP as control under doxycycline-inducible promoter. (Figure 5h, Figure S10a/b/c, Supplementary Table 7). The analysis of the resulting AP-MS data confirms that each of the three purifications enriches the other two complex members compared to the control. This in turn confirms that interactions between the three proteins are not mutually exclusive, which is also in agreement with published binary interaction data obtained by other methods annotated on BioGRID^13,19,38^. Overall, all our data support a model where PTPN14 promotes the interaction between LATS1/2 and YAP1, the subsequent LATS1/2 mediated phosphorylation of YAP1 which, in turn, leads to YAP1 inactivation and cytoplasmic retention (Figure 5i).

## DISCUSSION

Intra- and intercellular signaling systems largely depend on the modulation of the cellular proteome at different levels, including alteration of protein abundance, modification and interactome remodeling. Most proteomic measurements of signaling systems to date have focused on the exhaustive analysis of single proteomic layers, exemplified by the analysis of altered protein abundance profiles^44^ and/or the analysis of altered phosphorylation patterns^45^. Yet, it is well-known that molecular events at the different layers are interdependent and collectively determine the state of the signaling system^9,46,47^. An integrated view of the reorganization of the proteome across layers in the context of the cellular state is therefore critically important to unravel the underlying signaling network. In this study, we developed a generic experimental and computational approach to study the interdependence between PPIs and phosphorylation in the context of signaling systems. Using the Hippo signaling system as a model, we combined genetic and chemical perturbation with protein and phospho-protein correlation profiling following complex fractionation by BNPAGE, to identify different YAP1 proteoforms associated with distinct complexes, recapitulate known mechanisms of regulation and provide new insights into the PTPN14-mediated inactivation of YAP1.

In the first step of the study, we used two different classes of phosphatase inhibitors for in depth profiling of YAP1 phosphorylation states and YAP1 protein interaction dynamics. The combined data evidenced how changes in YAP1 phosphorylation correlated with significant changes in the interactome; and how proteins known to be functionally related – e.g. TEAD proteins, apicobasal proteins, and the F-box proteins BTRC and FBXW11 - undergo coordinated changes. We then combined BNPAGE separation combined with MS analysis to isolated distinct YAP1 complexes and obtain a more granular map of the phosphosite/PPI relationship. This approach is limited by a number of important factors, including accidental co-migration of non-interacting proteins; the interpretation-neutral application of a signal process algorithm, which may miss important features revealed only by manual inspection; and the correlative (as opposed to causal) relationship that can be established between phosphosites and complexes. In spite of these limitations, our approach increases greatly the depth as compared to previous co-fractionation experiments, which have been (but for a few exceptions) ^24^ carried out in lysates^5,6,48,49^; it has indeed proven capable of isolating complexes reported in literature (i.e. module 1 contains the complex RICH1/AMOT polarity complex^26^) or whose members are otherwise functionally related; and on top assigned distinct YAP1 proteoforms to each of the 9 identified complexes. By this means, our data reduces the number of potential YAP1 proteoform-complexes associations by several orders of magnitude and paves the way for establishing causal relationships. In this sense, we also consider our approach complementary to peptide-based pulldowns^8^, which establishes direct relationships between single peptides and interacting partners, but can neither describe complexes nor multi-phosphosite dependencies of PPIs.

Finally, we monitored changes in the phosphorylation status and interactome of YAP1 in a panel of cell lines lacking known Hippo regulators. The results confirm the role of LATS1/2 and NF2 as main modulators of YAP1 function and support the role of PTPN14 as an additional critical negative regulator of YAP1 transcriptional activity as demonstrated in several earlier studies and in different cell systems^34,38,39, 50–53^ . We verified by immunoblotting, immunofluorescence and proteomics that LATS1/2 and PTPN14 KOs affect the activity and localization of YAP1 in similar ways, albeit at different magnitudes. Our targeted proteomics approach revealed a distinct pattern of hyperphosphorylated YAP1 sites in PTPN14 KO cells that closely matches the one measured in LATS1/2 mutants and also showed that PTPN14 is required for the interaction of YAP1 with LATS1/2, highlighting the benefits of coordinately measuring changes in PPI and phosphorylation patterns. Furthermore, we show by reciprocal quantitative AP-MS that PTPN14, YAP1 and LATS1 are binding to each other in a non-mutually exclusive fashion, indicating the existence of a trimeric complex. The existence of this PTPN14-YAP1-LATS1 complex was also apparent from our AP-BNPAGE data, showing their distinct co-migration as part of the nuclear module (module 3, Figure 3b/c). Taken together these data suggests a model where PTPN14 may supports LATS dependent YAP1 phosphorylation via trimeric complex formation. It has been shown that increasing cell density and the extent of cell-cell contacts which is accompanied by a strengthened interaction of YAP1 with LATS and PTPN14, also leads to an augmented phosphorylation of YAP1^35,54^. It is tempting to speculate that PTPN14, by supporting the formation of YAP1-LATS complex, may be a key player in enabling the cell density-dependent YAP1 interactome reshaping and signaling.

In summary, we describe a strategy to simultaneously analyze two critical aspects of cell signaling – complex formation and phosphorylation – as well as their interdependence combining multiple layers of proteomics data. Besides representing a comprehensive and sensitive account of YAP1 PTMs and interactors, our data suggest a model for PTPN14 as a negative YAP1 regulator by supporting the LATS binding and phosphorylation of YAP1. Given the widespread nature of phosphorylation controlled complex formation, we strongly believe that the presented strategy represents a significant analytical advance to disentangle regulatory mechanisms for a wide range of cellular signaling systems.

## MATHERIAL AND METHODS

### Plasmids and cloning

Expression constructs were generated with a N terminal Strep-HA-tagged bait proteins and entry clones of a Gateway compatible human clone collection (ORFeome v5.1 and v8.1). The integration of the entry clones into the Gateway destination vectors (pcDNA5/FRT/TO/SH/GW)^55^ was performed with an enzymatic LR clonase reaction (Invitrogen).

### Tissue culture and DNA transfection

T-REx^TM^ Flp-In cell lines purchased from Invitrogen were cultured in DMEM (4.5 g/l glucose, 2 mM L-glutamine) (Gibco), supplemented with 10% fetal bovine serum (FBS) (BioConcept), 100 U/ml penicillin (Gibco) and 100 µg/ml streptomycin (Gibco). HEK293A cell lines were purchased from Invitrogen or received as gift by the Guan lab^30^ were cultured in DMEM (4.5 g/l glucose, 2 mM L-glutamine), supplemented with 10% FBS (BioConcept), 100 U/ml penicillin (Gibco), 100 µg/ml streptomycin (Gibco) and MEM Non-Essential Amino Acids Solution. Cell lines were cultured at 37 °C in a humidified incubator with 5% CO2.

### Stable cell line generation of N terminal Strep-HA-tagged proteins

T-REx^TM^ Flp-In cells were co-transfected with the corresponding expression plasmid and the pOG44 vector (Invitrogen) encoding the Flp-recombinase using jetPrime (Polyplus) according to the manufacturer’s instructions. Two days after the transfection, cells were selected in hygromycin (100 µg/ml) and blasticidin C (15 µg/ml) containing medium for 3 weeks.

### CRISPR/Cas9-mediated gene knock-out of PTPN14 in HEK293A cells

To generate CRISPR/Cas-9 PTPN14 KO cells we designed guideRNAs based on their specificity score from the Optimized CRISPR Design web tool (http://crispr.mit.edu) (PTPN14 gRNA target sequence 1: 5’ – CACCGCGTTGTAGCGCCGTGTCCGGCGG (exon 1) , PTPN14 gRNA target sequence 2: 5’ – CACCGGCTCCACCCATCGTGCTTGCTGG (exon 2)). Annealed DNA oligonucleotides containing the target sequence were cloned into the hSpCas9 plasmid (pX458, Addgene) using BbsI restriction sites. Subsequently, HEK293A cells were transfected with two hspCas9 constructs encoding gRNAs with the target sequence 1 and 2. The cell culture medium was replaced 4 hours after transfection and cells were recovered for 72 hours. Then, 1×10e6 cells were gently detached from the tissue culture plate with 0.25% trypsin-EDTA (Gibco) and resuspended in PBS containing 1% FBS. GFP-expressing cells were detected and isolated by FACS (BD Facs Aria IIIu sorter) and sorted into a 96-well plate. The cell clones were expanded and characterized by western blotting and mass spectrometry.

### Western Blot

Cells were grown in 6 well plates to 80% confluency and harvested. Cell pellet was snap frozen and lysed in 100 µl lysis buffer (0.5% NP40, 50 mM HEPES (pH 7.5), 150 mM NaCl, 50 mM NaF, 400 nM Na_3_VO_4_, 1mM PMSF and protease inhibitor cocktail). The cell lysate was cleared by centrifugation (15000g for 20 min), boiled for 5 min after addition of 3X Laemmli sample buffer, loaded on NuPAGE 4-12% Bis-Tris SDS-PAGE gels (Invitrogen) for gel electrophoresis and then transferred onto nitrocellulose membranes (Trans-Blot Turbo, BioRad). The following primary antibodies were used: anti-PTPN14 (#13808, Cell Signaling), anti-actin (#179467, Abcam), anti-YAP1 (#15407, Santa Cruz), anti-YAP1phosphoS127 (#4911, Cell Signaling), anti-HA (HA.11,901513, BioLegend). Proteins were detected by enhanced chemiluminescence (ECL, Amersham) using horseradish-peroxidase-coupled secondary antibodies (Rabbit #7074, Cell Signaling and Mouse #115035003, Jackson ImmunoResearch).

### Protein extraction and full proteome digestion

Cells were cultured in 150 mm tissue culture plates untill they reach 80% confluence. Cells were harvested and the cell pellet was snap frozen and lysed. Lysis was performed in 8 M urea and subjected to harsh sonication (3 times 1 minute, 80% amplitude and 80% cycle time, Hielscher-Ultrasound Technology), Benzonase (Sigma) activity (50U/ml) and centrifugation (14000g for 15 minutes). The protein amount of the cleared supernatant was measured by the Bicinchoninic acid (BCA) assay (Pierce) and 50 µg protein were subsequently reduced (5 mM TCEP in 50 mM ammonium bicarbonate, 30 min) and alkylated (10 mM iodoacetamide, 30 min). The protein sample was diluted to 1.5 M urea and proteolyzed with 0.5 μg of LysC (Wako) and 2 μg Trypsin (Promega, sequencing grade) for 16 h at 37 °C. Proteolysis was quenched by 0.1% TFA and peptides were purified with a C18 column (Sep-Pak 1cc, Waters). Eluted peptides were dried using a speed vacuum centrifuge before being resuspended in 20 μl 0.1% formic acid and 2% acetonitrile. iRT peptides (Biognosys) were spiked to each sample (1:50) before LC-MS/MS analysis for quality control.

### Affinity purification of SH tagged proteins and digestion (AP-MS)

The expression of SH-tagged bait proteins stably integrated in T-REx^TM^ Flp-In cells was induced with 1 µg/ml tetracycline for 24 h. For affinity purification three or four (based on bait expression), 150 mm tissue culture plates at 80% cell confluency were harvested and the cell pellet was snap frozen. The frozen pellet was lysed with the following buffer (HNN lysis buffer): 0.5% NP40, 50 mM HEPES (pH 7.5), 150 mM NaCl, 50 mM NaF, 400 nM Na_3_VO_4_ supplemented with 1mM PMSF, 1.2 µM Avidin (IBA) and protease inhibitor cocktail (P8849, Sigma), using 800 µl of lysis buffer for each lysed cell plate. The lysate was incubated on ice for 20 min and subjected to mild sonication (3 times 10 seconds, 35% amplitude and 80% cycle time, Hielscher-Ultrasound Technology) and digestion of nucleic acids via Benzonase (Sigma) (50 U/ml). The cleared cell lysate was incubated with 50µl crosslinked Strep-Tactin Sepharose beads (IBA) for 1 h on a rotation shaker. Before the incubation with lysate, beads were crosslinked with 5 mM of di-succinimidylsuberate DSS (Thermo) in 50 mM HEPES (pH 8.0), 150 mM NaCl for 30 minutes at 37 °C with strong agitation and quenched with 50 mM ammonium bicarbonate for 30 minutes at 37 °C. Upon washing two times with lysis buffer and three times with HNN buffer (50 mM HEPES (pH 7.5), 150 mM NaCl, 50 mM NaF), beads and bound proteins were transferred in 10 kDa molecular weight cut-off spin column (Vivaspin 500, Sartorious), following the FASP protocol^56^. Briefly, beads in solution were centrifuged at 8000g until dryness. Samples were denatured, reduced (8 M Urea and 5 mM TCEP in 50 mM ammonium bicarbonate, 30 min) and alkylated (10 mM iodoacetamide, 30 min). Each sample was subsequently washed three times by flushing the filter with 25 mM ammonium bicarbonate and digested with 0.5 μg of Trypsin (Promega, sequencing grade) for 16 h at 37 °C. Proteolysis was quenched by 0.1% TFA and peptides were purified with a C18 microspin column (Nest Group). Eluted peptides were dried using a speed vacuum before being resuspended in 20 μl 0.1% formic acid and 2% acetonitrile. For quality control, iRT peptides (Biognosys) were spiked to each sample (1:50) before LC-MS/MS analysis. In fractionated samples, peptides were subjected to high pH fractionation in reversed phase (microspin column, Nest Group) following the procedure based on the high pH reversed-phase peptide fraction kit (Pierce).

### In vivo treatment of YAP1 SH tagged with phosphatase inhibitors

The expression of YAP1 N terminal SH-tagged integrated in T-REx^TM^ Flp-In cells was induced with 1 µg/ml tetracycline. After 24 hours, media was exchanged with growth media and cells were stimulated with 100 μM and 150nM of Vanadate and Okadaic acid (Biovision) for 2 or 20 minutes, and 60 or 150 minutes, respectively. Pervanadate was freshly prepared by mixing on ice for 20 minutes Na_3_VO_4_ (Sigma Aldrich) with H_2_O_2_ in a molar ratio 1:5, following the protocol of Huyer et al.^57^. After stimulation, cells were harvested and the cell pellet was snap frozen.

### AP-BNPAGE of YAP1 complexes (AP-BNPAGE-MS)

A visualized and detailed description of the protocol to resolve purified protein complexes is published by Pardo et al.^58^ The experimental procedure described below underlines the important and critical steps to perform the experiment. For affinity purification coupled with Blue Native separation, fifteen 150 mm tissue culture plates at 80% cell confluency, treated with 1 µg/ml tetracycline for 24 h were harvested and the cell pellet was snap frozen. Cells were lysed, cleared and incubated with 50 µl of Strep-Tactin Sepharose beads, following the conditions described above for the affinity purification of SH tagged proteins and digestion (AP-MS). Upon washing two times with lysis buffer and three times with HNN buffer (50 mM HEPES (pH 7.5), 150 mM NaCl, 50 mM NaF), bound proteins were incubated for 30 minutes and eluted with 50 µl of 2.5 mM biotin in HNN buffer (Thermo). 40 µl of eluted protein was supplemented with 12 µl of native sample loading buffer and loaded on Native PAGE 3-12% Bis Tris precast protein gels (Invitrogen) for native separation, according to the manufacturer’s instruction. Different from instructions, the cathode chamber was only filled with Light Blue Cathode Buffer. Native PAGE gel separation was performed for 3 hours at 4 °C applying three step gradient voltage (150V-180V-200V). Once the separation was finished, proteins were stained with Simple Blue Safe Stain (Invitrogen) and proteolyzed following Protease MAX Surfactant (Promega) in gel digestion protocol. To excise 64 protein bands with the same size from a native gel separation (necessary for quantitative proteomics data), a custom device constituted by 100 parallel blades spaced 1 mm from one another was used. Briefly, protein bands were distained, shrunk, reduced (25 mM DTT) and alkylated (55 mm iodoacetamide) before proteolysis. Digestion was performed in 50 µl digestion solution (0.5 µg of Trypsin (Promega, sequencing grade), 0.1 µg of LysC (Wako), 0.01% ProteaseMAX Surfactant (Promega) in 50 mM ammonium bicarbonate). After overnight digestions, peptides extracted in solution were collected, while gel pieces were covered with 50% acetonitrile solution for 30 minutes to improve the yield of the peptide extraction. Peptide solutions generated from the proteolysis and from the treatment of gel pieces with 50% acetonitrile solution were dried and resuspended in 10 μl 0.1% formic acid and 2% acetonitrile. iRT peptides (Biognosys) were spiked to each sample (1:50) before LC-MS/MS analysis for quality control.

### Immuno-affinity purification using custom-designed YAP1 antibodies. Design of epitope and beads preparation for IP-MS

To perform antibody based purification, we designed a custom antibody against the C terminal region YAP1 (TLEGDGMNIEGEELM). The following parameter were determinant for the peptide choice: i) exposition and lack of secondary structure (we used Psipred^59^ as secondary structure prediction tool), ii) low sequence homology with other human proteins, iii) non-involvement of PTMs and protein interactions, iv) peptide stability in solution (we used ProtParam Tool from Expasy to monitor the stability). The peptide was synthetized, coupled to KLH carrier protein and used for rabbit immunization with the “*Speedy 28-Day program*” by Eurogentec. The final bleed was affinity purified in AKTA pure chromatography system (GE Healthcare) with the epitope antibody column with 50 mM HEPES (pH 7.5), 150 mM NaCl as running buffer and 0.1 M Glycine (pH=3) for the elution. The column for the affinity purification was prepared coupling the peptide TLEGDGMNIEGEELM to NHS group of HiTrap NHS-Activated affinity column (GE Healthcare). Eluate was neutralized in Tris base solution 100 mM, pH 8.8, dialyzed overnight in buffer 50 mM HEPES (pH 7.5), 150 mM NaCl using membrane dialysis tube (Pur-A-Lyzer Mega Dialysis 3500KDa) (Thermo). The dialyzed eluate and the flow through obtained from peptide affinity purification were quantified, affinity characterized and coupled to protein A Sepharose 4 Fast Flow (GE Healthcare) following the protocol^60^. Briefly, 10 mg of specific and un- specific antibodies were incubated with 5 ml of wet protein A Sepharose 4 Fast Flow beads for one hour, beads were extensively washed with 0.2 Sodium Borate pH=9 and crosslinked with 20 mM of DMP for one hour. After quenching reaction with ethanolamine 0.2 M, beads were aliquoted (∼200 µg of antibody per purification) and ready to use.

### Immuno-affinity purification using custom-designed YAP1 antibodies (IP-MS)

HEK293A and HEK293A with genetic deletions were cultured in ten 150 mm tissue culture plates to 80% confluency, harvested and the cell pellet was snap frozen. The frozen pellet was lysed in 8ml of lysis buffer: 0.5% NP40, 50 mM HEPES (pH 7.5), 150 mM NaCl, 50 mM NaF, 400 nM Na_3_VO_4_ supplemented with 1 mM PMSF and protease inhibitor cocktail (P8849, Sigma). The lysate was incubated on ice for 20 min and subjected to mild sonication (3 times 10 seconds, 35% amplitude and 80% cycle time, Hielscher-Ultrasound Technology) digestion of nucleic acids via Benzonase (Sigma) (50 U/ml). The cleared cell lysate was incubated with protein A beads coupled with antibodies overnight on a rotation shaker. After incubation, beads were washed and proteolyzed following the conditions described above for the affinity purification of SH tagged proteins and digestion (AP-MS).

### IF analysis

200,000 HEK 293A cells were seeded on poly-lysine coated glass coverslips and grown with the growth media as described above. After 24 hours, cells were washed in ice-chilled 1X PBS and fixed with 4% PFA. Permeabilizing with 0.1% Triton, cells were blocked with 5% filtered BSA containing 0.01% Triton for at least an hour. Cells were probed with anti-YAP1 primary antibody (Santa Cruz Biotechnology, sc-376830) at 1:100 dilution and alexa488-labeled secondary antibody at 1:2000 dilution. Before the final wash of coverslips with 1X PBS, cells were incubated with 1:3000 DAPI for 10 minutes in the dark. Subsequently, the slides were mounted onto glass slides and imaged using inverted Nikon Eclipse Ti microscope. The nuclear relocation of YAP1 was imaged at 63× oil objectives. The acquisition of images in relevant channels was controlled using open-source software *micromanager*. Z-stack of images at multiple positions were acquired using the piezo drive and automated XY drive.

Image analysis was conducted using CellProfiler software. Images in two channels- DAPI (nucleus) and Cy5 measuring YAP1 levels were imported into the CellProfiler. Prior to analysis, illumination function was calculated in both channels by selecting the background function, block size of 60, and “Fit Polynomial” smoothing method. The correction function was calculated based on all images in each channel and subsequently applied to the corresponding channel to obtain illumination corrected images. The corrected DAPI image was used to segment the nucleus and define the “Nucleus” as a primary object. Propagating from coordinates of Nucleus into corrected YAP1 signal in Cy5 channel using “Global” threshold strategy, a secondary object encompassing the whole cell was created. Subtracting the Nucleus object from thus propagated cell, a tertiary object called “Cytoplasm” was created. Furthermore, two objects were created, expanding 2 pixels and 10 pixels from the nucleus. Subsequently, a tertiary objected called “ring” was created around the nucleus by subtracting 2-pixel expanded nucleus from the 10-pixel expanded nucleus. This ring was further limited within the cells by masking it within the coordinates of “Cytoplasm” object, defining it as “Perinuclear”. Finally, the median intensity of corrected YAP1 signal (Cy5 channel) was measured within the Nucleus and Perinuclear objects and the ratio between the two was computed to determine relative nuclear relocation of YAP1. The experiment was repeated three independent times and more than 1300 single cells from three repeats were analyzed per condition. Student’s t-test was performed between single cell data from each condition to determine the statistical significance. The significance is indicated with *** for *P*<0.001.

### qPCR analysis

HEK293A cell lines (WT, LATS1/2KO and PTPN14KO) were grown in one 60mm dish at 50% confluence. Cells were detached by trypsinization and lysed using QIAshredder columns (Qiagen). Total RNA was extracted using RNeasy kit (Qiagen) and DNA was degraded using RNase-free DNase I (Qiagen) following manufacturer instructions. RNA was reverse transcribed into cDNA using random hexanucleotides (Microsynth) and SuperScript II polymerase (Roche). The relative abundance of CTGF and CYR61 mRNA was determined using a Roche LightCycler and SYBRgreen (Roche). GAPDH was used as a reference gene.

The following oligos were used:

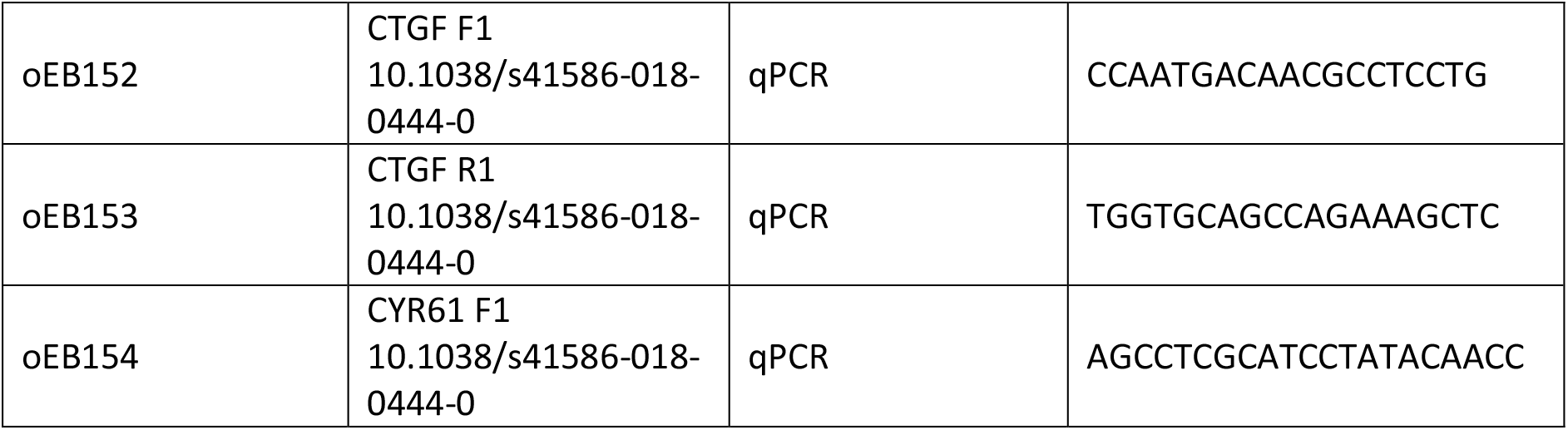
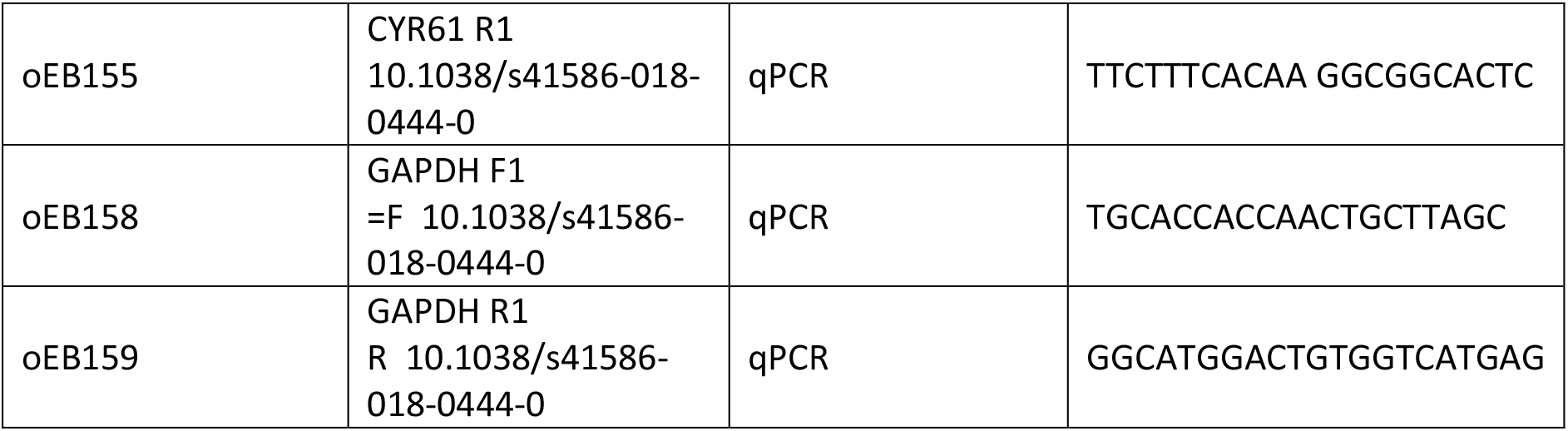

### Mass spectrometry based data acquisition

#### MS data acquisition of in vivo phosphatase treatment of YAP1 SH tagged

LC-MS/MS analysis was performed on an Orbitrap Elite mass spectrometer (Thermo Scientific) coupled to an Easy-nLC 1000 system (Thermo Scientific). Peptides were separated on a Thermo PepMap RSLC column (15 cm length, 75 µm inner diameter) with a 60 min gradient from 5% to 35% acetonitrile at a flow rate of 300 nl/min. The mass spectrometer was operated in data-dependent acquisition (DDA) mode with the following parameters: one full FTMS scan (350-1600 m/z) at 120’000 resolution followed by fifteen MS/MS scans in the Ion Trap. Charge states lower than two and higher than seven were rejected. Selected ions were isolated using a quadrupole mass filter of 2.0 m/z isolation window. Precursors with MS signal that exceeded a threshold of 500 were fragmented (CID, Normalized Collision Energy 35%). Selected ions were dynamical excluded for 30 s.

#### MS data acquisition of AP-BNPAGE of YAP1 complexes (AP-BNPAGE-MS)

LC-MS/MS analysis was performed on an Orbitrap Q Exactive HF mass spectrometer (Thermo Scientific), coupled to an Acquity UPLC M-class system (Waters). Peptides were loaded on commercial trap column (Symmetry C18, 100Å, 5µm, 180 µm*20mm, Waters) and separated on a commercial column (HSS T3, 100Å, 1.8µm, 75 µm*250mm, Waters) using a 40 min gradient from 8% to 30% acetonitrile at a flow rate of 300 nl/min. The mass spectrometer was operated in data-dependent acquisition (DDA) mode with the following parameters: one full FTMS scan (350-1500 m/z) at 60’000 resolution, 15 ms injection time and 3e6 AGC target, followed by 12 FTMS/MS scans at 60’000 resolution, 110ms injection time and 1e5 AGC target. Charge states lower than 2 and higher than 7 were rejected. Selected ions were isolated using a quadrupole mass filter of 1.2 m/z isolation window and fragmented (HCD, Normalized Collision Energy 28%). Selected ions were dynamical excluded for 20 s.

#### MS data acquisition for targeted analysis of genetic KO screen in HEK293A cell lines

LC-MS/MS analysis was performed on an Orbitrap Q Exactive HF mass spectrometer (Thermo Scientific) coupled to an Acquity UPLC M-class system (Waters). Peptides were loaded on commercial trap column (Symmetry C18, 100Å, 5µm, 180 µm*20mm, Waters) and separated on a commercial column (HSS T3, 100Å, 1.8µm, 75 µm*250mm, Waters) using a 90 min gradient from 5% to 35% acetonitrile at a flow rate of 300 nl/min. The mass spectrometer was operated in parallel reaction monitoring (PRM) mode with the following parameters: one full FTMS scan (400-1500 m/z) at 120’000 resolution, 250 ms injection time and 3e6 AGC target, followed by time scheduled target PRM scans at 120’000 resolution, 247 ms injection time and 2e5 AGC target. Selected ions were isolated using a quadrupole mass filter of 2.0 m/z isolation window and fragmented (HCD, Normalized Collision Energy 30%). Scan windows were set to 10 min for each peptide in the final PRM method. The inclusion list with targeted peptides analyzed is reported (Supplementary Table 3).

#### MS data acquisition of YAP1 IP-MS

LC-MS/MS analysis was performed on an Orbitrap Q Exactive HF mass spectrometer (Thermo Scientific) coupled to an Acquity UPLC M-class system (Waters). Peptides were loaded on commercial trap column (Symmetry C18, 100Å, 5µm, 180 µm*20mm, Waters) and separated on a commercial column (HSS T3, 100Å, 1.8µm, 75 µm*250mm, Waters) using a 60 min gradient from 2% to 37% acetonitrile at a flow rate of 300 nl/min. The mass spectrometer was operated in data-dependent acquisition (DDA) mode with the following parameters: one full FTMS scan (350-1500 m/z) at 60’000 resolution, 15 ms injection time and 3e6 AGC target, followed by twelve FTMS/MS scans at 60’000 resolution, 110 ms injection time and 5e4 AGC target. Charge states lower than two and higher than seven were rejected. Selected ions were isolated using a quadrupole mass filter of 1.2 m/z isolation window and fragmented (HCD, Normalized Collision Energy 28%). Selected ions were dynamical excluded for 30 s.

#### MS data acquisition of targeted analysis of YAP1 IP-MS in HEK293A cell lines

LC- MS/MS analysis was performed on an Orbitrap Fusion Lumos Tribrid mass spectrometer (Thermo Scientific) coupled to an EASY-nLC 1200 system (Thermo Scientific). Peptides were separated on Acclaim PepMap 100 C18 (25 cm length, 75 µm inner diameter) with a 90 min gradient from 5% to 35% acetonitrile at a flow rate of 300 nl/min. The mass spectrometer was operated parallel reaction monitoring (PRM) mode with the following parameters: one full FTMS scan (200-2000 m/z) at 30’000 resolution, 54 ms injection time and 1e6 AGC target, followed by time scheduled target PRM scans at variable resolution and injection time (15’000 R/22ms IT; 30’000 R/54ms IT; 60’000R/118ms IT; 120’000 R/246 ms IT). Selected ions were isolated using a quadrupole mass filter of 1.4 m/z isolation window and fragmented (HCD, Normalized Collision Energy 27%). Scan windows were set to 10 min for each peptide in the final PRM method. The inclusion list with target peptides analyzed is reported (Supplementary Table 5).

#### MS data acquisition of total protein expression in a panel of HEK293A cell lines with genetic deletion

Assay library generation: LC-MS/MS analysis was performed on an Orbitrap Fusion Lumos Tribrid mass spectrometer (Thermo Scientific) coupled to an EASY-nLC 1200 system (Thermo Scientific). Peptides were separated on Acclaim PepMap 100 C18 (25 cm length, 75 µm inner diameter) with a 120 min gradient from 3% to 35% acetonitrile at a flow rate of 300 nl/min. The mass spectrometer was operated in data-independent acquisition (DDA) mode with the following parameters: one full FTMS scan (350-2000 m/z) at 120’000 resolution (400 m/z), 50 ms injection time and 4e5 AGC target, followed by twelve FTMS/MS scans at 30’000 resolution (400 m/z), 54 ms injection time and 5e4 AGC target for a cycle time of 3 seconds. Charge states lower than two and higher than seven were rejected. Selected ions were isolated using a quadrupole mass filter of 1.4 m/z isolation window and fragmented (HCD, Normalized Collision Energy 35%). DIA. measurements: samples were analyzed with the same set up used for assay library generation. The mass spectrometer was operated in data-independent acquisition (DIA) mode with the following parameters: one full FTMS scan (375-1250 m/z) at 120’000 resolution, 50ms injection time and 4e5 AGC target, followed by 40 variable windows from 375 to 1250 m/z with 1 m/z overlap at 30’000 resolution, 54ms injection time and 1e5 AGC target for a cycle time of 3 seconds. Precursor ions were fragmented with HCD, Normalized Collision Energy 35%.

#### MS data acquisition of targeted analysis of PTPN14, YAP1, LATS1 and GFP SH-tagged AP-MS

LC-MS/MS analysis was performed on an Orbitrap Q Exactive HF mass spectrometer (Thermo Scientific) coupled to an Acquity UPLC M-class system (Waters). Peptides were loaded on commercial trap column (Symmetry C18, 100Å, 5µm, 180 µm*20mm, Waters) and separated on a commercial column (HSS T3, 100Å, 1.8µm, 75 µm*250mm, Waters) using a 60 min gradient from 5% to 35% acetonitrile at a flow rate of 300 nl/min. The mass spectrometer was operated parallel reaction monitoring (PRM) mode with the following parameters: one full FTMS scan (350-1800 m/z) at 60’000 resolution, 110 ms injection time and 1e6 AGC target, followed by time scheduled target PRM scans at 60’000 resolution, 119 ms injection time and 2e5 AGC target. Charge states lower than two and higher than seven were rejected. Selected ions were isolated using a quadrupole mass filter of 2.0 m/z isolation window and fragmented (HCD, Normalized Collision Energy 30%). Scan windows were set to 6 min for each peptide in the final PRM method. The inclusion list with target peptides analyzed is reported. (Supplementary Table 7).

### Experiment design, data process and statistical analysis of mass spectrometry data

#### Analysis of in vivo treatment of YAP1 SH tagged with phosphatase inhibitors

The experiment was performed with three independent biological replicates of YAP1 SH tagged purification without stimulation and with vanadate stimulation (2 and 20 minutes) or with okadaic acid stimulation (60 and 150 minutes). To identify YAP1 interactors, we analyzed twelve purification controls with GFP SH tagged. Acquired spectra were searched using the MaxQuant software package version 1.5.2.8 embedded with the Andromeda search engine^61^ against human proteome reference dataset (http://www.uniprot.org/, downloaded on 10.10.18) extended with reverse decoy sequences. The search parameters were set to include only full tryptic peptides, maximum one missed cleavage, carbamidomethyl as static peptide modification, oxidation (M) and phosphorylation (S, T, Y) as variable modification and “match between runs” option. The MS and MS/MS mass tolerance were set, respectively, to 4.5 ppm and 0.5 Da. False discovery rate of <1% was used at the protein level to infer the protein presence. The protein abundance was determined from the intensity of top two unique peptides for each protein. Interactome definition: high confident interactors of AP-MS experiments were determined by SAINTexpress^20^ with default parameters using spectral counts obtained from Max Quant analysis (MS/MS Count). Twelve SH-GFP pulldowns processed and measured in parallel with the samples and additional control runs from the CRAPome database (http://crapome.org/62) were used to filter high confidence interactors of YAP1 (SAINT threshold score > 0.90). MS1 quantification of phosphorylated peptides: phosphorylated peptides were filtered based on Andromeda phospho localization probability score (>0.8). Furthermore, phospho-sites that were not detected in all three replicates in at least one condition were filtered out. Phospho- peptide intensities were bait normalized and missing value were imputed with the median of biological replicates (only one missing value per replicate per condition was allowed. MS1 quantification of interactors: LFQ protein intensities of high confidence interactors were bait normalized and missing values were imputed with the median of biological replicates (only one missing value per condition) or using random sampling from a normal distribution generated 5% less intense values. Two sided t test and *p* (corrected for multiple hypothesis testing using the Benjamini-Hochberg method) were computed to compare treated and control groups. Cluster of kinetic profiles for interactors was performed with normalization to unstimulated samples and with a fuzzy cluster algorithm (mfuzzy package, R).

#### Analysis of AP-BNPAGE of YAP1 complexes

The experiment was performed with single analysis of YAP1 SH tagged purification, the eluate was separated with blue native gel and fractionated in 64 protein bands.

Identification of YAP1 interactors: three independent biological replicates of YAP1 SH tagged purification were proteolyzed and peptides were fractionated using High pH Reversed-Phase Fractionation Kit. Protein identified in fractionation samples were filtered using SAINT express, as described above, to obtain a deeper list of high confidence interactors (57 proteins). Proteins in the list of YAP1 interactors were considered for the AP-BNPAGE experiment.

In the AP-BNPAGE-MS experiment, acquired spectra were searched using the MaxQuant software package using specification described above in the analysis of in vivo treatment of YAP1 SH tagged with phosphatase inhibitors. Protein intensities of YAP1 interactors (high confidence interactors list from fractionated YAP1 interactome, Figure S2h) and phosphosite intensity of YAP1 and YAP1 interactors were extracted from the protein and peptide matrices. LFQ protein intensity was normalized using iRT peptide intensity; phospho peptides were filtered based on phospho localization probability score (filter peptides with score above 0.8 in at least one fraction; filter fractions with score above0.5) and the intensity was normalized using YAP1 protein abundance (only YAP1 phospho- peptides) and for iRT peptide intensity. Missing values were imputed with the average of two neighboring fractions. Phospho-peptide and protein profiles were normalized for the maximum value across the fractionation dimension. Next, each profile was split based on identified peaks using gaussian smoothing function (minimum normalized intensity 0.2 and width 2 for proteins; minimum normalized intensity 0.3 and width 2 for phospho- peptides). In the analysis of interactors, YAP1 was excluded as the protein is identified in all fractions and interacts with all protein groups identified in the separation. Hierarchical clustering based on the distance of peak correlation was performed for interactors and phospho-sites to generate co-migration groups. The number of clusters and the cluster stability was evaluated by the Silhouette plot using Euclidian distance of clusters.

Recall rate for protein-protein interactions (PPIs) identified with observed cluster was calculated for each comigration group with the ratio between identified protein-protein interactions and protein-protein interactions annotated in BioGRID (version 3.5.176)^13^. In the ratio calculation of BioGRID annotated interactions, YAP1 interactions were not considered. GO cellular component enrichment (https://david.ncifcrf.gov/63) was calculated with the ratio between proteins involved in “cell junctions”, “cytoplasm”, “cytosol”, “apical plasma membrane”, “nucleus cell compartments”, over all proteins identified for each comigration group (24 proteins). Phospho cluster enrichment, was calculated with the ratio between proteins involved in the three clusters described in figure 2E after vanadate and okadaic acid treatment (cluster 1,2,3), over all proteins identified for each comigration group (24 proteins). For PPIs recall rate, GO CC enrichment and phospho cluster enrichment analysis random clusters were generated from 1000 random clusters generated from YAP1 identified interactors (24 proteins), including always YAP1 and with the same group size of the observed clusters. Two generated distributions were assayed for normality with Shapiro test and with a two sided t-test for difference. The layout of protein-protein interaction comigration groups (Figures 3c.) was generated using Cytoscape (v3.6.0)^64^.

#### Analysis of targeted quantification of genetic KO screen in HEK293A cell lines

The experiment was performed in three independent biological replicates. Supplementary Table 3 reports the list of all target peptides and proteins measured in the analysis. PRM assay containing protein knockout in the cell line panel, housekeeping protein (Actin B) and iRT peptides was generated from spectra library data imported in Skyline (v.4.1)^65^. Spectra libraries were built using published spectral libraries^66^ and Mascot search results (v. 2.4.1, MatrixScience) after proteomic analysis of cell lysates and YAP1 affinity purified as described above. Briefly, for Mascot search with precursor tolerance of 15 ppm and fragment tolerance of 0.6 Da, a Mascot score larger than 20 and an expectation value smaller than 0.05 were considered to identify correctly assigned peptides. Peak group identification and automatic peak picking of six fragment per peptide was performed employing the mProphet ^67^ algorithm. The second best peaks were used as controls for the training model. For peptide identification we used the following criteria: retention time matching to spectra library within 5% of the gradient length and dot product between library spectra intensities and light peptides > 0.75. After identification, peptide abundance was obtained from the sum of the integrated area of three fragment ions per peptide. Fragment ions with a signal to noise ratio less than 5 were filtered out for the quantification. Peptide values were normalized for the intensity of housekeeping peptides (Actin B) and for the intensity of iRT peptides.

#### Analysis of YAP1 IP-MS

he experiment was designed with YAP1 immuno-purification and two different control purifications (co-immuno purification with unspecific antibodies in HEK293A wt cells and anti-YAP1 co-immunopurification in YAP1KO HEK293A cells). All purifications were performed in three independent biological replicates. All samples were fractionated with reverse phase high pH fractionation kit (Pierce). Acquired spectra were searched using the MaxQuant software package using specification described above in the analysis of in vivo treatment of YAP1 SH tagged with phosphatase inhibitors. Proteins significative upregulated (*p* value with Benjamini and Hochberg method correction < 0.05) in YAP1 immunopurifications with both purifications were considered as YAP1 interactors.

#### Targeted YAP1 IP-MS analysis in HEK293A cell lines

The experiment was performed with three independent biological replicates of YAP1 endogenous immune-purified from a panel of cell lysates.

For the targeted assay/panel, selected peptides belong to proteins, which were prior characterized within this study as high confidence interactors (identified in AP-MS and IP-MS experiments) were considered. This targeted panel was supplemented with YAP1 phosphopeptides (identified in AP-MS and IP-MS experiments). Supplementary Table 5 reports the list of all target peptides and proteins measured in the analysis. Isotope-labeled heavy peptides corresponding to the proteotypic peptides selected for this study, and containing either heavy lysine (13C(6) 15N(2)) or arginine (13C(6) 15N(4)) residues were purchased from JPT Peptide Technologies GmbH. Peptides were analyzed manually, and correct identification with six fragment ions per peptide was assigned based on the coelution of light and heavy peptide and matching peak shape for precursor and product ions from light and heavy peptides. The abundance of peptides was analyzed by summing the integrated areas of three fragment ions per peptide. Fragment ions with a signal to noise ratio less than 5 were filtered out for the quantification. Peptide intensity values were normalized for the intensity of 8 YAP1 peptides, for the TIC and for the intensity of iRT peptides. Significance of change in intensity was estimated with *p* values using two-sided, not paired t-test.

#### Analysis of total protein expression in a panel of HEK293A cell lines with genetic deletion

Differential protein expression of three independent biological replicates was measured by data independent acquisition (DIA). For library generation a single cell lysate from HEK293A was proteolyzed and peptides were fractionated and analyzed in DDA mode. Hybrid spectral library was generated by Spectronaut 13 (version 13.2.190709)^68^ (Biognosys) using peptide identified in the DIA runs and peptides identified in DDA mode from a prior offline fractionation (8 fractions, reverse phase high pH fractionation). For DDA analysis, acquired spectra were searched using the MaxQuant software package using specification described above in the analysis of in vivo treatment of YAP1 SH tagged with phosphatase inhibitors, excluding threonine, tyrosine and serine phosphorylation as variable modification. The generated library included entries for 120’844 peptide precursors and 8’408 protein groups. For the DIA analysis, extraction of quantitative data was performed with Spectronaut querying the library above-mentioned with the following settings: tolerance of 10 ppm for precursor and 25 ppm for fragment ions and a dynamic retention time extraction window with nonlinear iRT retention time calibration. Precursor and proteins were identified with q value cut-off of 0.01 (5,947 protein groups). Data normalization by total ion current (TIC)) and filtering was performed with mapDIA^69^, where a standard deviation factor of 2 and a minimal correlation of 0.2 were used to filter robust fragment ions with minimum intensity threshold of 200. Filter strategy at protein level include minimum identification of two peptides per protein group in at least two of the three biological replicates. Group comparison level between each of the 7 conditions against the wildtype HEK293A signal was performed within mapDIA. We identified 31295 and 4436 proteins with only 3% of missing values across the matrix. Missing values were imputed with the median value of biological replicates (only one missing value per condition) or using random sampling from a normal distribution generated 1% less intense values. ANOVA statistical test was performed to compare protein profiles in all different cell lines.

#### Analysis of targeted PTPN14, YAP1, LATS1 and GFP SH-tagged AP-MS

The experiment was performed with three independent biological replicates of YAP1, PTPN14, LATS1, GFP SH tagged purifications. Supplementary Table 7 reports the list of all target peptides. Selected peptides from the YAP1 library were identified and quantified with the same criteria described above in the analysis of targeted YAP1 IP-MS in HEK293A cell lines. Peptide intensity values were normalized for the intensity of a reference peptide in the SH-tag of all bait proteins (AADITSLYK) and for the intensity of iRT peptides^70^.

## ACKNOLEDGEMENTS

We thank Dr. A. Leitner for technical mass spectrometry support; Dr. N. Selevsek and Dr. B. Roschitzki from the Functional Genomic Center Zürich for the access and maintenance of the mass spectrometer facility. We thank Dr. S.W. Plouffe and Dr. K.L. Guan for providing the panel of HEK293A cell lines with genetic deletions of YAP1 regulators. This work was supported financially by: IMI project ULTRA-DD (FP07/2007-2013, grant no.115766, F.U.,M.M.,F.F.,M.G.), Marie Curie Individual Fellowship (Grant agreement number 703759, R.C.), Innovative Medicines Initiative 2 Joint Undertaking (JU) under grant agreement No 875510 (M.G.), ETH Career Seed Grant (SEED-18 17-2, R.C.), long-Term Fellowship from the European Molecular Biology Organization (ALTF 928-2014, M.M.) and by European Research Council AdvG grant 670821 (RA).

## CONTRIBUTION

F.U., R.C. and M.G. conceived the study. F.U., R.M., F.F. E.S.B. and M.M. performed the experiments. F.U., R.C., A.F. analyzed the data. A.F. and F.F. contributed to the experimental design. N.T. and M.P. provided valuable input for experiments. R.C., F.U., M.G., R.A. drafted the manuscript, which has been revised and approved by all authors. R.A. and M.G. supervised the project and provided funding.

## LEGENDS FOR TABLES S1, S2, S3, S4, S5, S6, S7

**Supplementary Table 1**. Results from analysis of in vivo treatment of YAP1 SH tagged with phosphatase inhibitors (Figure 2) (raw files list, MaxQuant output and statistical evaluation).

**Supplementary Table 2.** Results from analysis of AP-BNPAGE of YAP1 complexes (raw files list, MaxQuant output and statistical evaluation) (Figure 3).

**Supplementary Table 3.** Results from analysis of targeted analysis of genetic KO screen in HEK293A cell lines (raw files list, monitored peptides, skyline transitions output) (Figure 4).

**Supplementary Table 4.** Results from analysis of YAP1 IP-MS (raw files list, MaxQuant output, statistical evaluation) (Figure 4).

**Supplementary Table 5.** Results from analysis of targeted YAP1 IP-MS in HEK293A cell lines (YAP1 interactors and phosphosites) (raw files list, monitored peptides, skyline transitions output, statistical evaluation) (Figure 4).

**Supplementary Table 6.** Results from analysis of total protein expression in a panel of HEK293A cell lines with genetic deletion (raw files list, mapDIA input, and statistical evaluation) (Figure 5).

**Supplementary Table 7** Results from analysis of targeted PTPN14, YAP1, LATS1 and GFP SH-tagged AP-MS (raw files list, monitored peptides, skyline transitions output and statistical evaluation) (Figure 5).

---

**Figure S1.**
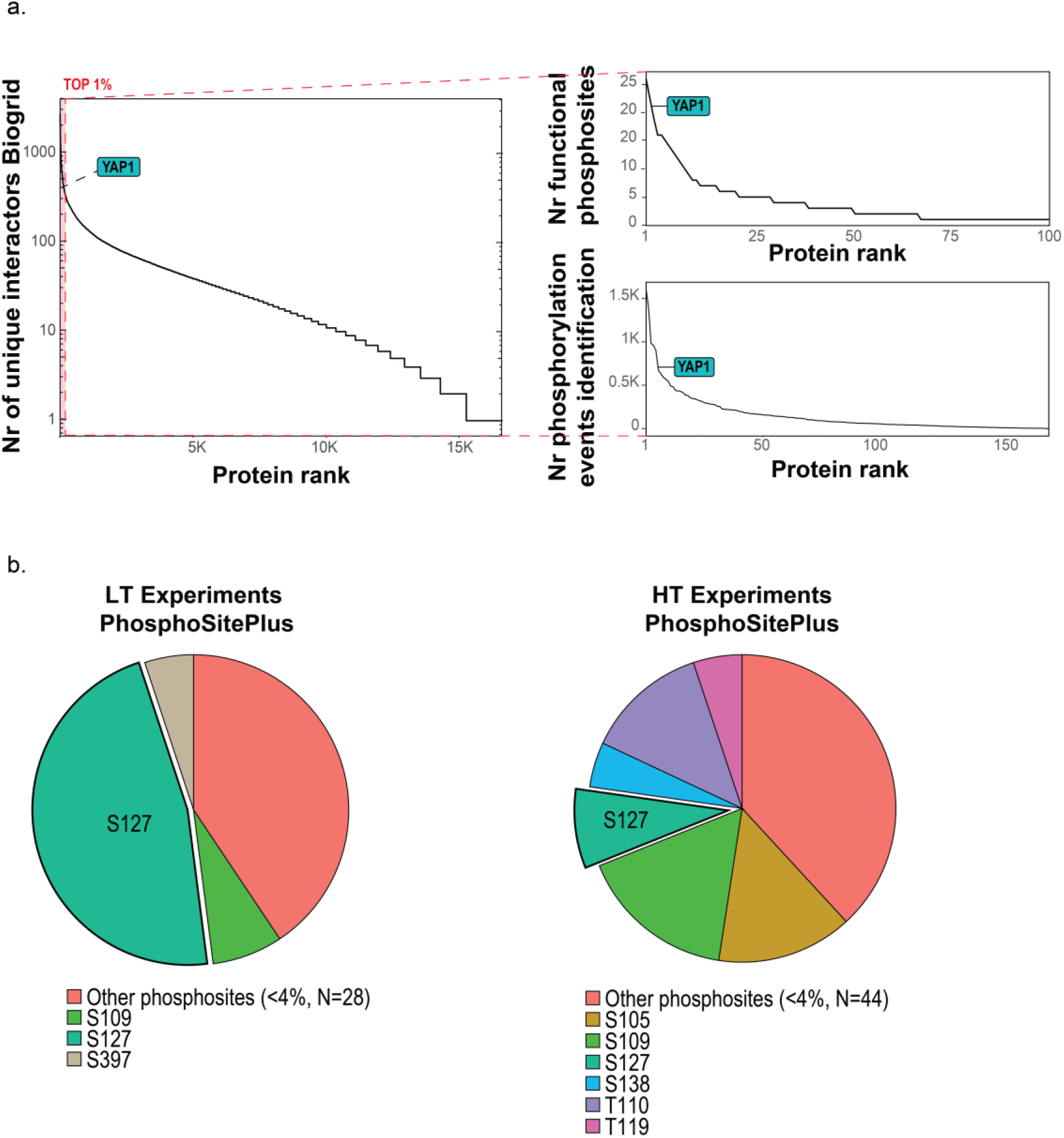
**a.** Selection of YAP1 as a model system was based on the number of known interactors (annotated in *BioGRID*; left) and number of identified and functional phosphosites (based on *Phosphositesplus* (CST) data and the scoring system proposed in Ochoa et al., 2019, respectively). **b.** Fraction of experiments (low throughput, left and high throughput, right) in which YAP1Phosphosites are annotated (*Phosphositesplus*, CST).

**Figure S2.**
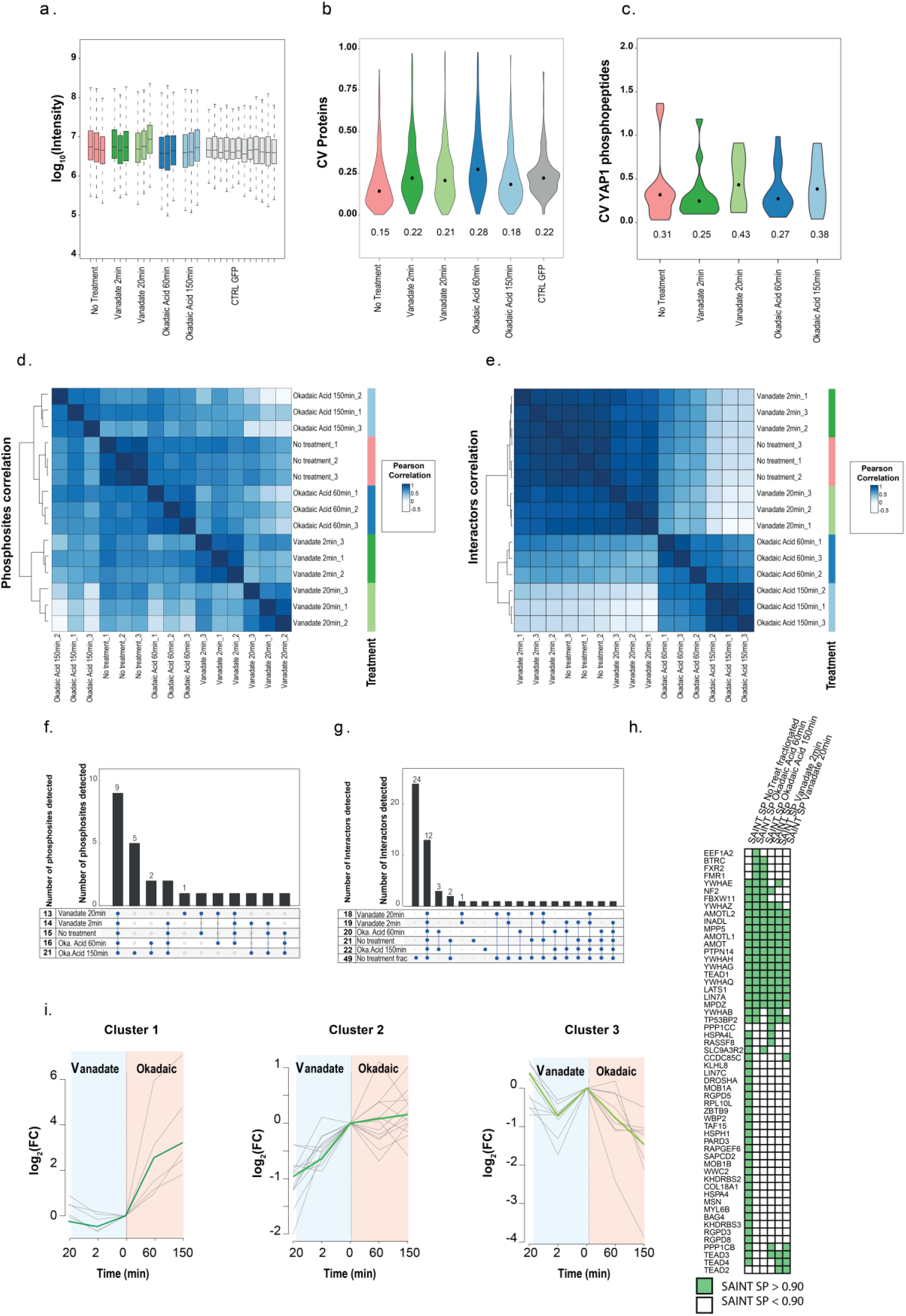
**a.** Distribution intensity (log10) of identified proteins. **b.** Distribution of coefficient of variation (CV) values at protein level. **c.** Distribution of coefficient of variation (CV) values for YAP1 phosphosites. **d.** Unsupervised hierarchical cluster based on Pearson correlation for YAP1 phosphosites identified across the tested conditions. **e.** Unsupervised hierarchical cluster based on Pearson correlation between YAP1 interactors identified across the tested conditions. **f.** Upset plot of size and overlap of phosphosite sets identified across the tested conditions. **g.** Upset plot of size and overlap of interactor sets identified across the tested conditions. **h.** Heatmap of YAP1 high confidence interactors identified using a SAINT SP score threshold of 0.90 in the tested condition.

**Figure S3.**
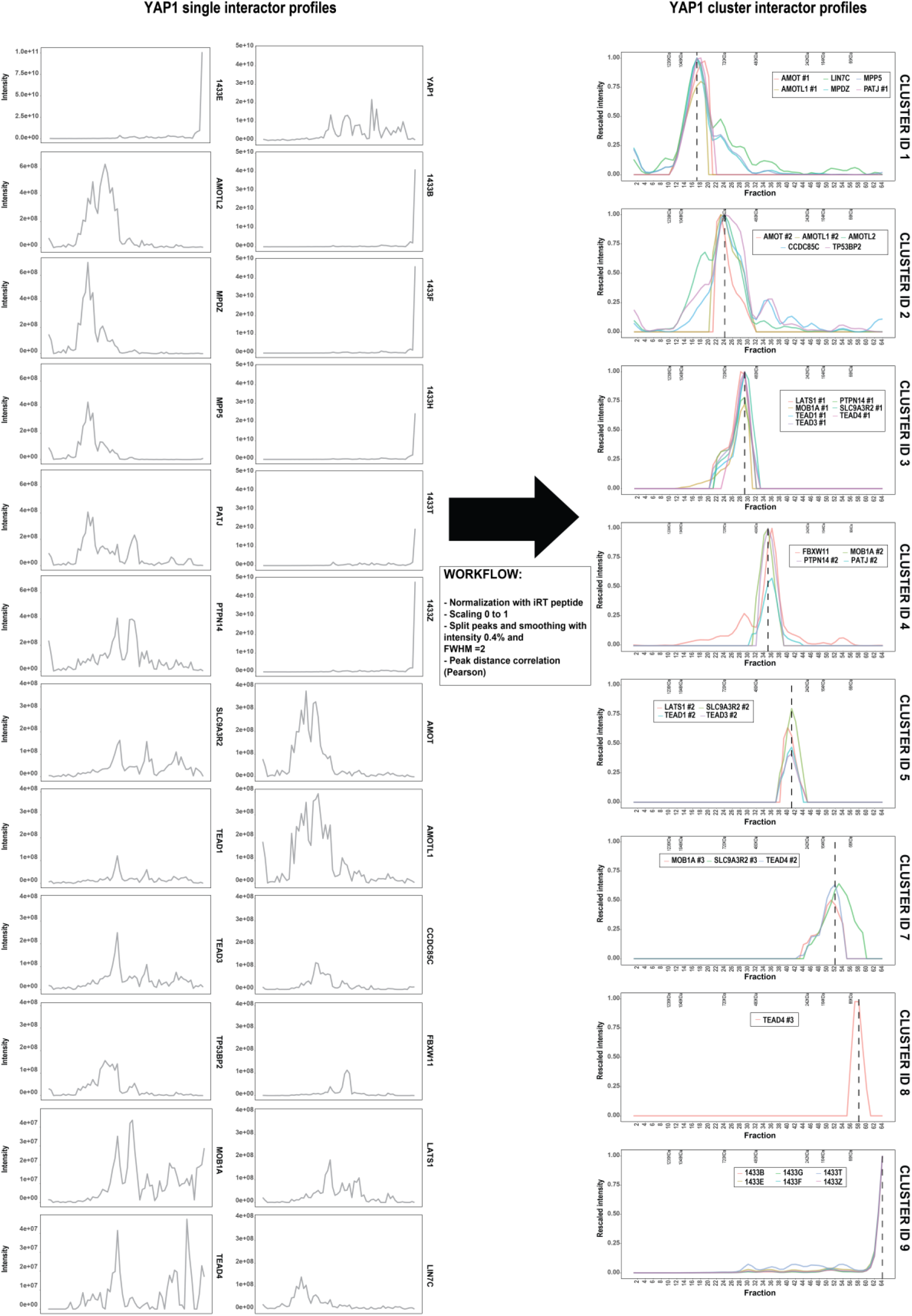
Raw intensity of AP-BNPAGE profiles of YAP1 interactors (left) and profiles after processing grouped by cluster membership (right).

**Figure S4.**
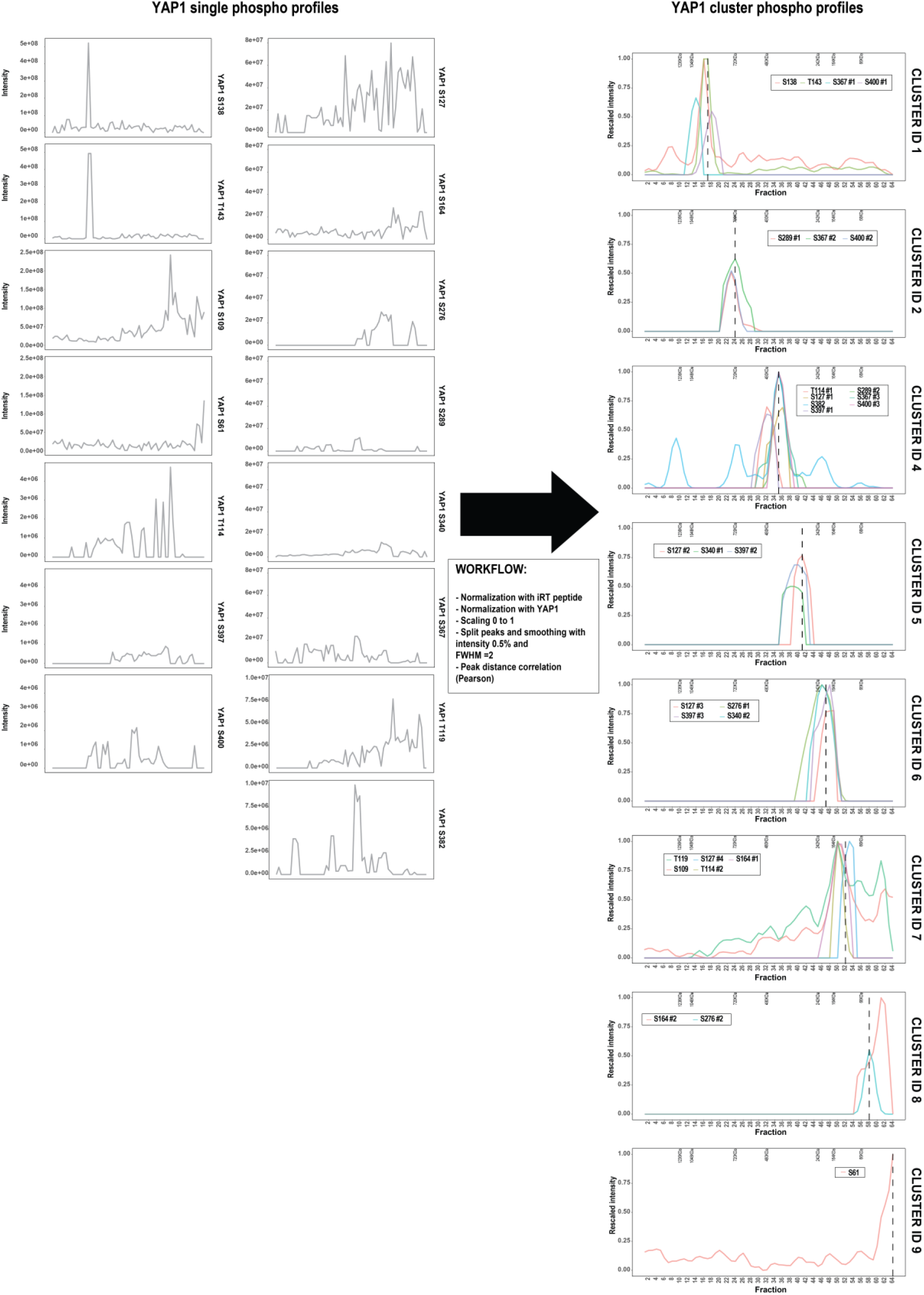
Raw intensity of AP-BNPAGE profiles of YAP1 phosphosites (left) and profiles after processing grouped by cluster membership (right).

**Figure S5.**
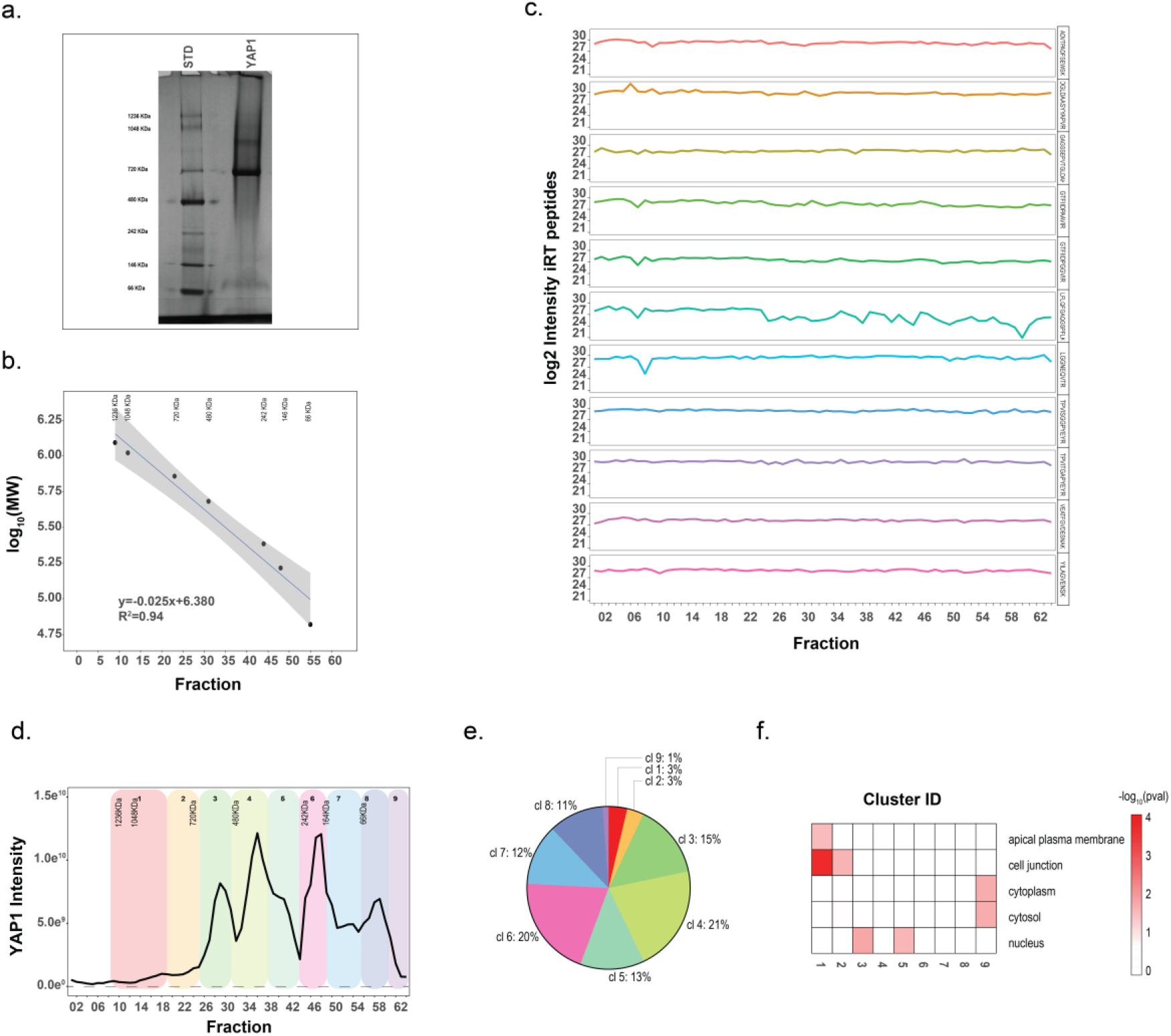
**a.** Coomassie staining of the BNPAGE used to resolve YAP1 complexes. **b.** Calibration curve with external standards (BNPAGE proteins standard) separated on BNPAGE. Calibration curve was used to estimate the molecular weight of YAP1 modules. **c.** Quantitative value of external standard (iRT peptides) spiked over all 64 fractions. Quantitative values are obtained from the integration of MS1 signal intensity. **d.** Distribution of smoothed signal of YAP1 MS1 intensity across 64 BNPAGE fractions. **e.** Relative YAP1 intensity associated with each of the identified clusters. **f.** Cellular component terms (GO) enriched in the identified modules indicates discrete protein localization.

**Figure S6.**
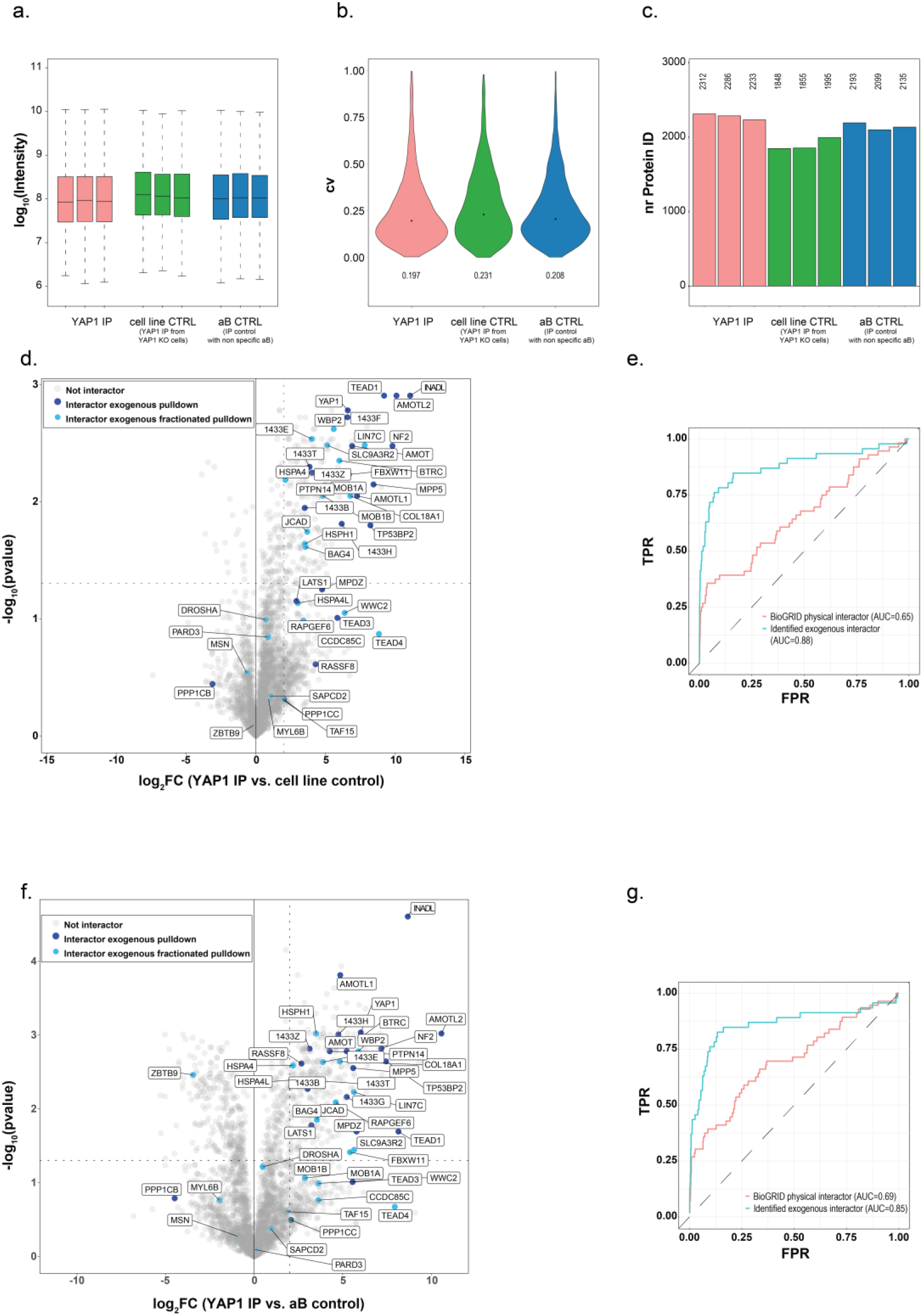
YAP1 endogenous interactome identified by YAP1 immuno-affinity purification. **a.** MS1 Intensity distribution (log10) of proteins identified in the indicated purifications (left: YAP1 IP-MS, center: cell line control, YAP1 IP-MS from YAP1 KO cells and right: aB control, non-specific control antibody IP-MS from HEK293). **b.** Distribution of coefficient of variation (CV) values of proteins identified in the indicated purifications. **c.** Number of proteins identified in the indicated purifications. **d**.**/f.** Volcano Plot of purified protein intensity (MS1) from YAP1 immuno-affinity purification and controls (cell line control **d.**; aB control **f.**). Protein identified and filtered as interactor (SP>0.9) in fractionated (light blue) and not fractionated (blue) AP-MS from HEK293 cells expressing epitope tagged YAP1 are annotated. **e/g.** Receiver-operating characteristic (ROC) curve and corresponding area under the curve (AUC) showing performance of YAP1 immune- affinity purification (using YAP1 KO cell line (**e.**) and aB control a (**g.**) controls) as benchmarked against YAP1 ectopically expressed APMS (blue line) and BioGRID annotated interactors.

**Figure S7.**
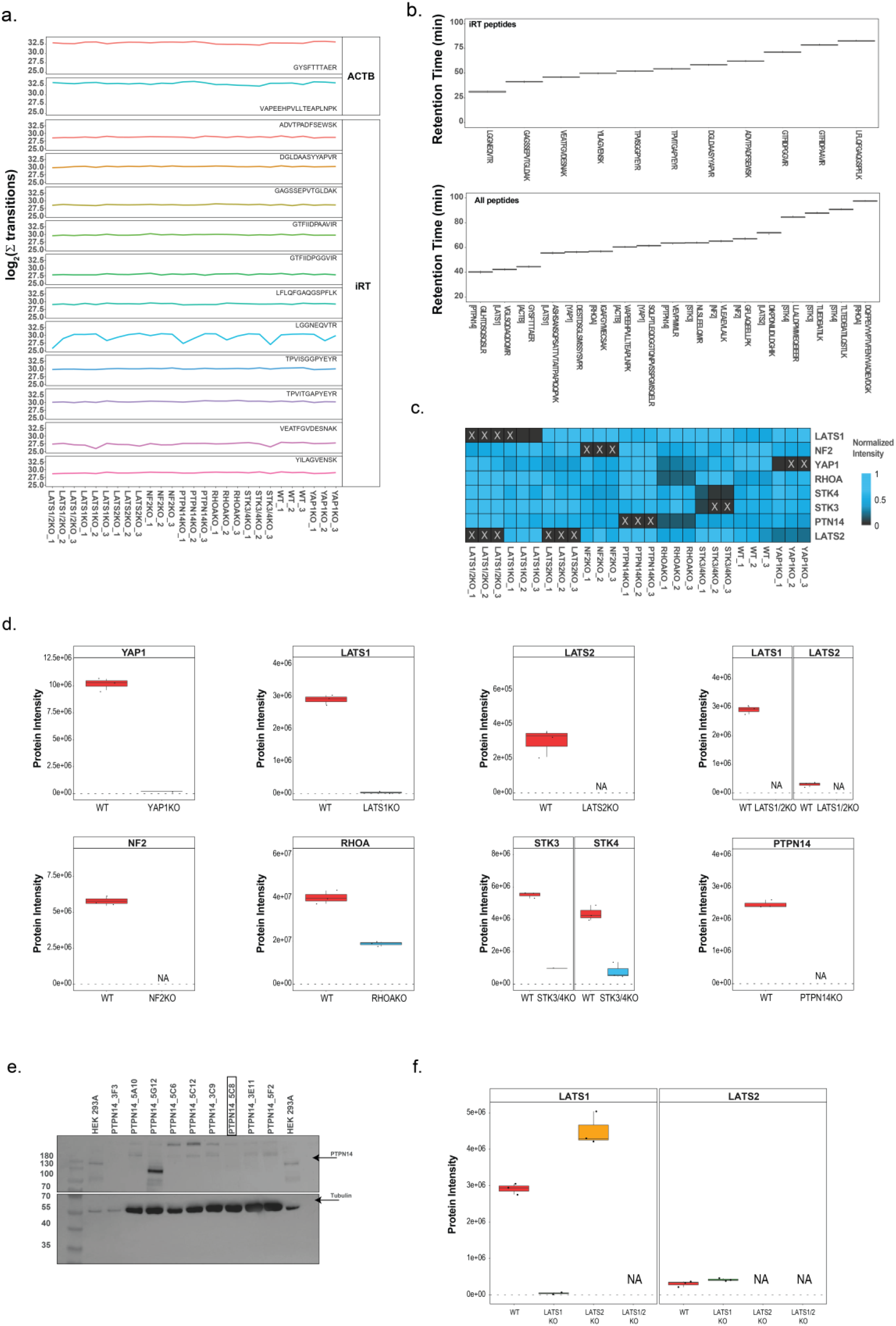
Targeted proteomic quantification and characterization of genetic deletion in nine different cell lines. **a.** Quantitative values of external standard (iRT peptides) spiked in the measurement for the targeted proteomic characterization of genetic deletions. Two peptides from Actin are used as loading control to normalize the injected lysate amount. Quantitative values are obtained from the sum of the transition values. **b.** Retention time of iRT (top) and selected peptides (bottom) used to evaluate genetic deletion efficiency. **c.** Characterization of genetic deletions in nine different cell lines. The heatmap reports the mean value of proteins intensities from three independent biological replicates normalized for the maximum detected value. **d.** Intensities of indicated proteins in the parental HEK293A control cell line (left, red) and the respective KO cell line (right, blue). **e.** Validation of PTPN14 deletion cell lines. Expression levels of PTPN14 in parental HEK293A control cell lines and CRISPR/Cas9 engineered PTPN14 KO clones in HEK293A cell lines as measured by Western blotting with the indicated antibodies. Clone PTPN14 5C8 has been selected for further analysis. **f.** Protein intensity abundance from three independent biological replicates of LATS1 (left) and LATS2 (right) in the indicated KO cell lines.

**Figure S8.**
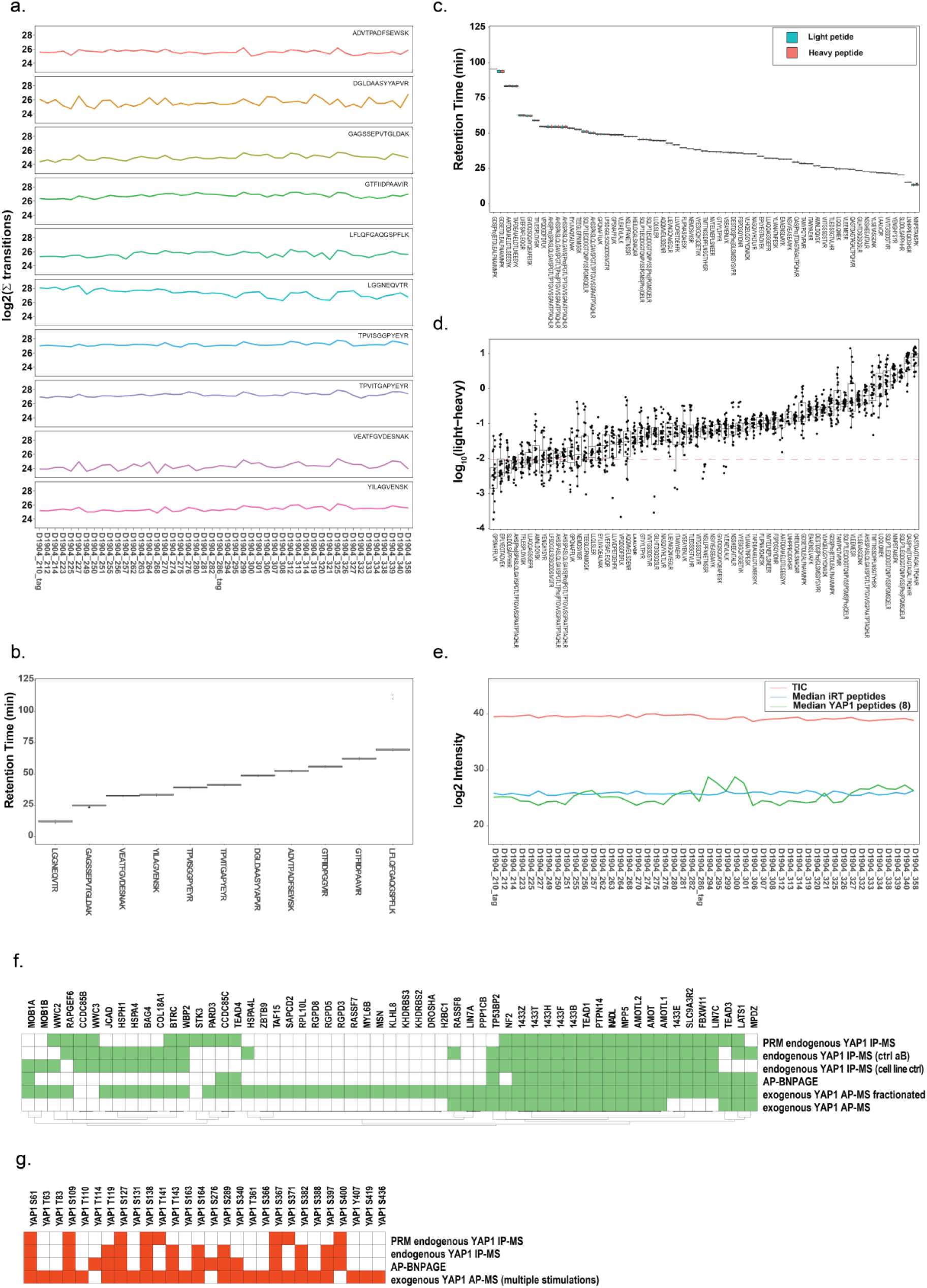
Targeted proteomic profiling of endogenous YAP1 phosphopeptides and interactors in cell lacking Hippo pathway members **a.** Intensities of external peptide standard (iRT peptides) spiked in the measurement for the targeted proteomic profile of YAP1 phosphopeptides and interactors in a panel of seven cell lines with Hippo genetic deletions. Quantitative values are obtained from the sum of the transition values. **b.** Retention time of iRT peptides. **c.** Retention time of monitored endogenous and reference peptides (light and heavy) for YAP1 interactors and phosphosites. **d.** Normalized intensity of monitored peptides expressed as the log10(light-heavy) peptide. All peptides are normalized using spiked in corresponding heavy reference peptides. **e.** Data are normalized based on TIC (Total Ion Current), median intensity of iRT peptides and mean intensity of YAP1 non phosphorylated peptides (8). All values used for the normalization are reported in the plot. **f.** Heatmap depicts all proteins monitored (identified and quantified with different approaches) across indicated experimental setups used in this study. **g.** Heatmap depicting all YAP1 phosphosites (identified and quantified with different approaches) in all different experiments setup used in this study.

**Figure S9.**
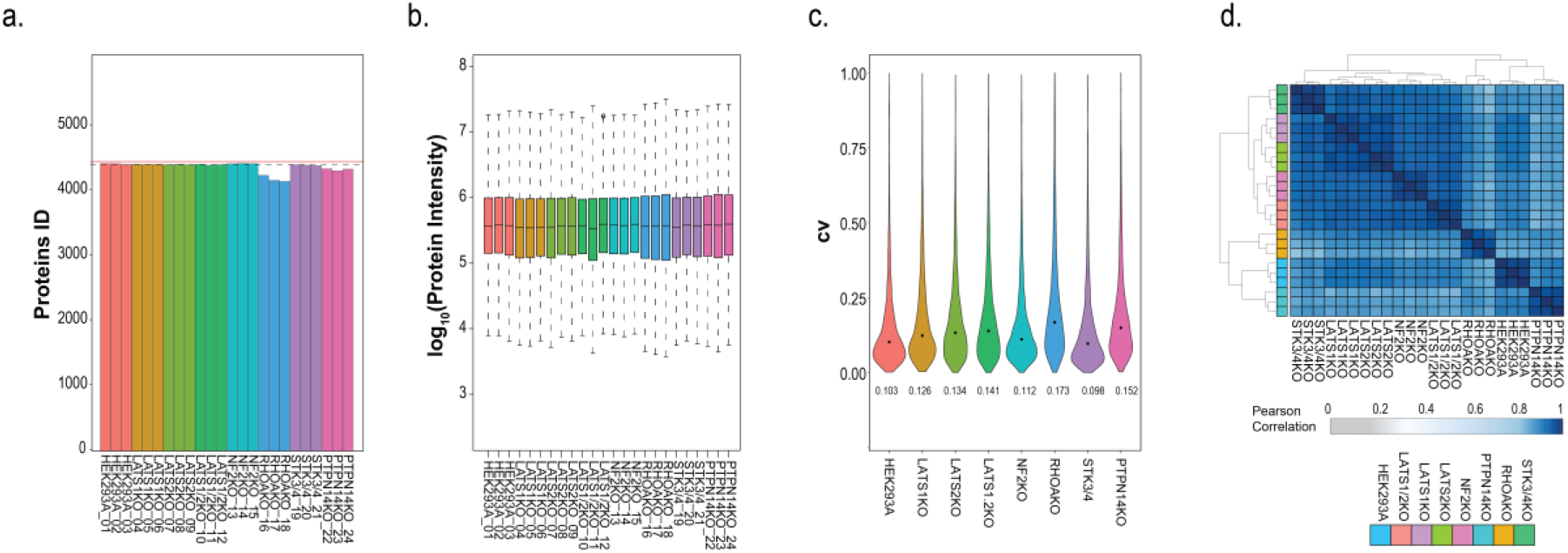
Differential protein expression data as determined by DIA proteomic workflow. **a.** Number of identified proteins in the DIA dataset. **b.** Distribution of protein intensity (log10) in the DIA dataset. **c.** Distribution of coefficient of variation (CV) values. **d.** Correlation matrix of protein intensities across indicated genetic backgrounds.

**Figure S10.**
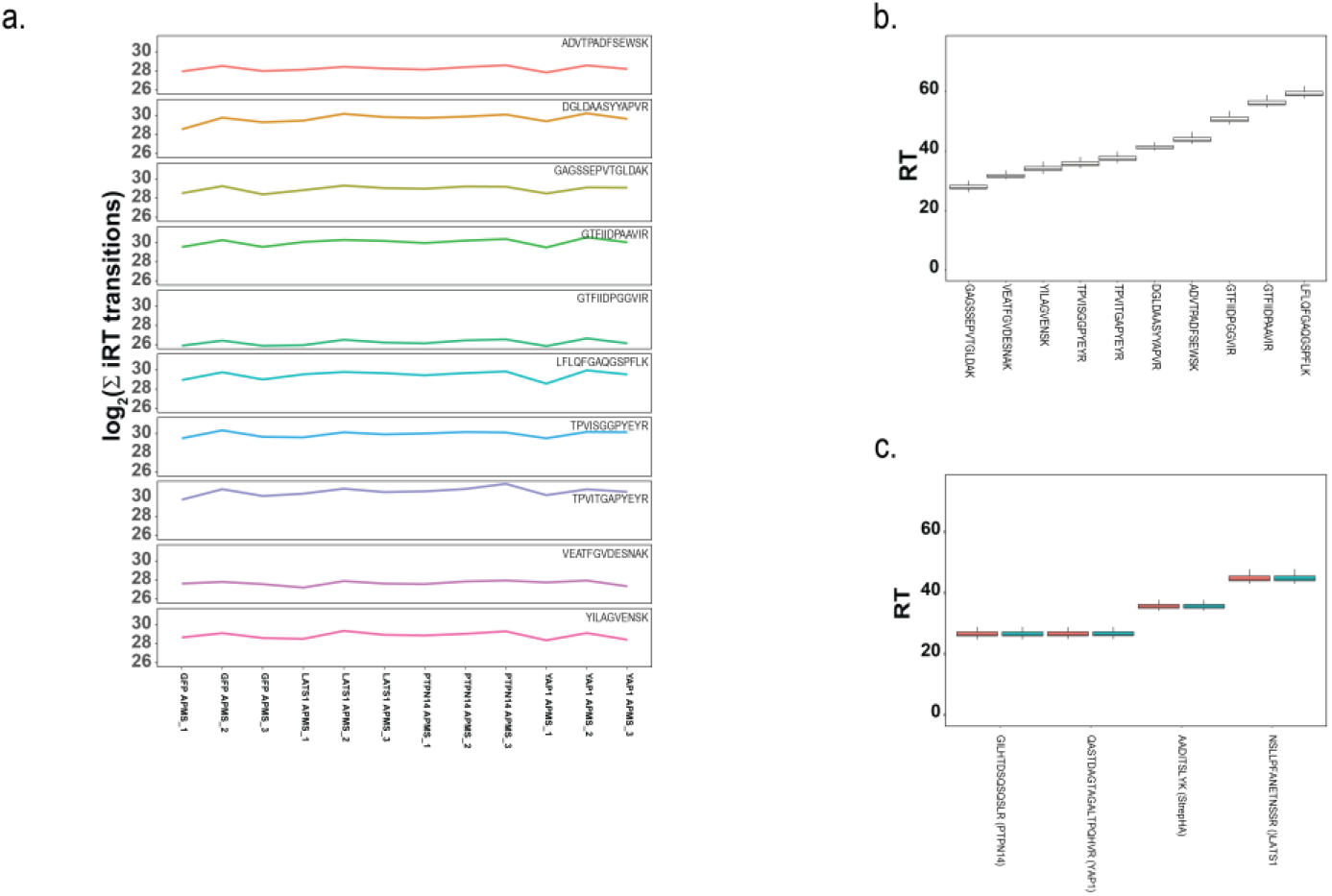
Targeted proteomic quantification of PTPN14, LATS1 and YAP1 in reciprocal AP-MS. **a.** Intensities of external peptide standard (iRT peptides) spiked in the measurement across different APMS experiments. Quantitative values are obtained from the sum of the transition values. Retention time of iRT peptides (**b.**) and monitored peptides (**c.**) used in the AP-MS experiment.

